# Spatially varying graph estimation for spatial transcriptomics cancer data

**DOI:** 10.1101/2025.05.04.652097

**Authors:** Satwik Acharyya, Jian Kang, Veerabhadran Baladandayuthapani

## Abstract

Modern spatial transcriptomic profiling techniques facilitate spatially resolved, high-dimensional assessment of cellular gene transcription across the tumor domain. The characterization of spatially varying gene networks enables the discovery of heterogeneous regulatory patterns and biological mechanisms underlying cancer etiology. We propose a *spatial Graphical Regression* (`sGR`) model to infer spatially varying graphs for high-resolution multivariate spatial data. Unlike existing graphical models, `sGR` explicitly incorporates spatial information to infer non-linear conditional dependencies through Gaussian processes. It conducts sparse estimation and selection of spatially varying edges, at both spatial and sub-spatial levels. Extensive simulation studies illustrate the profitability of `sGR` for spatial graph structural recovery and estimation accuracy. Our methods are motivated by and applied to two spatial transcriptomics data sets in breast and prostate cancer, to investigate spatially varying gene connectivity patterns across the tumor micro-environment. Our findings reveal several novel spatial interactions between genes related to immune activation and carcinogenesis regulation such as CD19 in breast cancer and ARHGAP family in prostate cancer. We also provide a modular software package for fitting and visualization of spatially varying graphs.

## 1 Introduction

### Scientific motivation

Spatial biology plays a crucial role in understanding the tumor microenvironment (TME) and is of immense importance in cancer research. Broadly, the TME is a complex cellular ecosystem consisting of various cell types, signaling molecules, extra-cellular components, and spatial organization [Loi et al., 2023]. Spatial profiling techniques enable researchers to investigate the spatial heterogeneity, interactions, and distributions of these components within the TME, providing critical insights into tumor development and progression [Yuan, 2016]. Techniques such as spatial transcriptomics (ST) based on in-situ hybridization, and multiplex immunofluorescence facilitate the assessment, characterization and visualization of the molecular spatial heterogeneity within the TME. While traditional high-throughput techniques such as bulk or single cell sequencing provide valuable information about tumor-specific gene expression programs; however, they fail to capture the spatial contextualization of these programs within the TME [Chen et al., 2023]. ST is a spatial genomics technology which combines single-cell RNA sequencing with spatial information of cells and it allows the mapping of gene expression patterns to specific locations within the tissue [Moses and Pachter, 2022]. The functionality of each cell is highly dependent on the interactions with its neighbours, measuring spatially-resolved transcription represents a crucial advancement in our ability to understand the spatial tissue structure and spatially varying interactions among genes. Spatial gene expression pattern helps to decipher complex signaling pathways, neighbourhood delineation and cell-cell interactions within a tissue [Saviano et al., 2020]. This enables researchers to study the spatial heterogeneity at a cellular level and characterize the underlying cancer etiology, for mechanism discovery and therapeutic potential.

Recent advancements of computational methods for ST data have been focused to identify spatially variable genes [Sun et al., 2020], spatial clustering analysis [Zhao et al., 2021], and spatially aware dimension reduction [Shang and Zhou, 2022]. However the study of spatially varying gene-gene association and transcriptional network modeling for ST data have received much less attention relative to the afore-mentioned methods. The dependency pattern between genes are closely regulated by networks of transcription factors and a biological understanding of these spatially varying networks can reveal novel disparities between the micro-environments of different tissue types [Pratapa et al., 2020]. A gene network is typically characterized by a graph where nodes denote the genes (or any genomic features) and edges represent the underlying dependency structure between genes. In case of ST data, these edges are spatially varying and the research questions are two-fold: (a) identification of spatially varying network structure between genes; and subsequently (b) explicit quantification of spatially varying dependency pattern both globally (over the whole spatial domain) and locally (at a sub-regional level) – enabling delineation of spatial patterns of gene-gene regulation in the TME at a systems level.

### Statistical formulation and inferential problem

A conceptual schema is provided in Figure 1A, where we have a tissue section image from prostate cancer (used in our application) which has been assayed for spatial gene expressions from *p* gene and *n* spatial locations. We show the plot of a gene expression in the middle panel and the resulting gene expression matrix is of *n* × *p* dimensions along with *n* × 2 spatial coordinates (shown in the last panel). Our goal is to estimate spatially varying graphs as shown in Figure 1B. The high-dimensional data structure – both in number of spatial locations (10^4^) and genes (10^2^) – creates the need for a flexible yet parsimonious model to estimate *O*(*np*^2^) elements of the spatially varying graphs.

**Figure 1.**
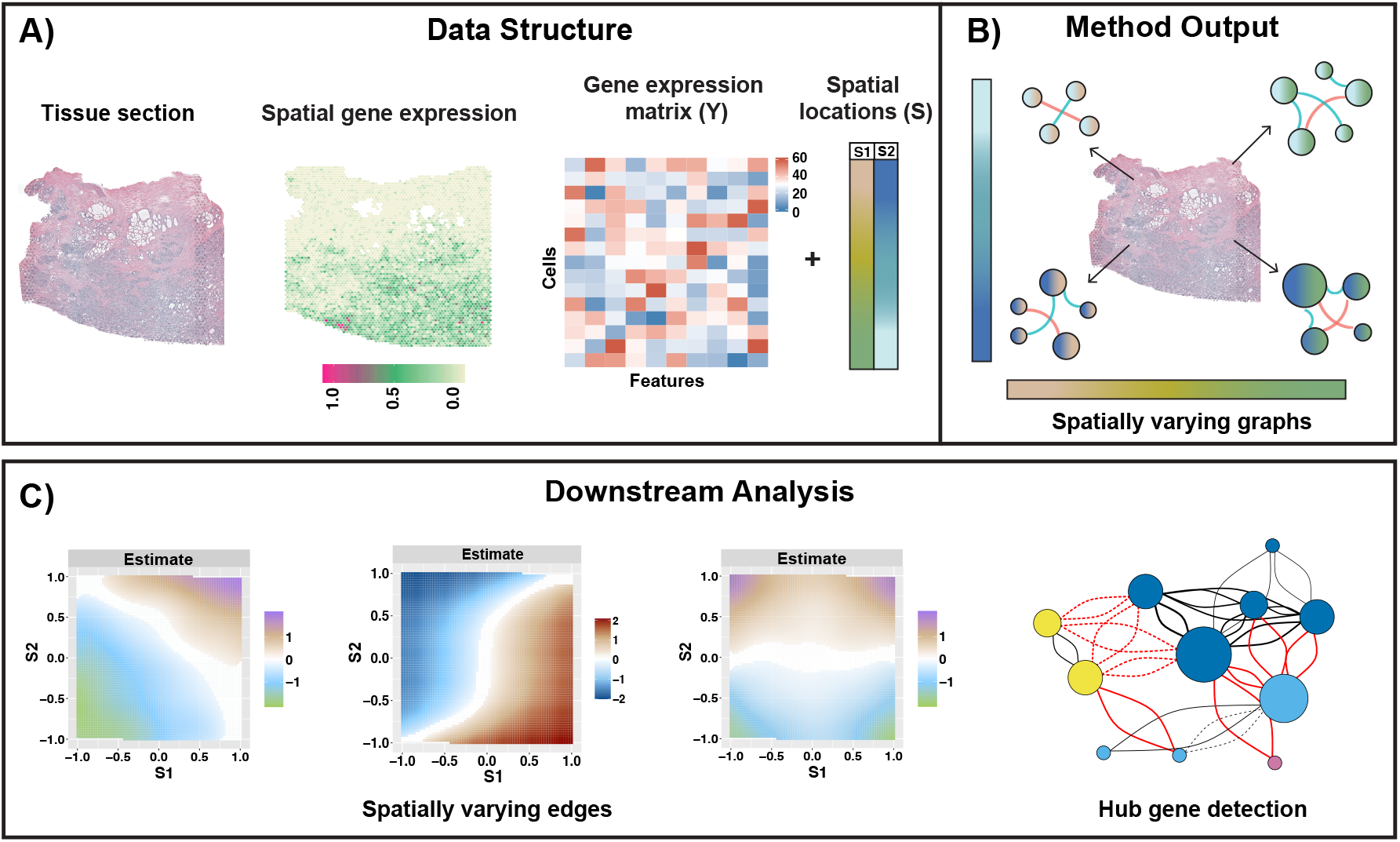
Overview of spatial graphical regression model: Left hand side of Panel A) shows the hematoxylin and eosin staining image (obtained from 10X genomics website) of the tissue of interest. Using spatial single-cell sequencing techniques, gene expression matrix and spatial locations are recorded from that tissue section as shown in right hand side of Panel A). We apply our spatial regression method on this data structure to obtain a spatial location specific graph which is spatially varying over the tissue domain as represented in Panel B). In Panel C), we have the spatially varying edges and connectivity degree based hub genes as a part of the downstream analysis from the spatially varying graphs in Panel B).

Graphical models are widely used methodology to study interactions which are quantified by conditional dependence (i.e. precision matrix) between genomic features in a complex biological system. Our main inferential goals are (I) characterization of graph topology changing over spatial domain, (II) identification of significant global and local spatial domains for spatially varying edges. The resulting spatial gene expression matrix in Figure 1A serve as an input to our framework which outputs a spatially varying network at each location over the spatial domain of the tissue section as displayed in Figure 1B. These networks can then be mined to infer spatial connectivity patterns over the tissue domain and identification of hub genes based on the (high) connectivity degree of specific genes (Figure 1C) for downstream biological interpretations – enabling discovery of biologically conserved transcription factors across different spatial regions.

### Existing methods and limitations

Gaussian graphical models (GGMs) are a standard class of probabilistic approach to analyzing conditional dependence structures, allowing assessment of the association between two random variables while factoring in the influence of all other variables [Lauritzen, 1996]. Solving the graph learning problem in GGMs entails both estimating the precision matrix and identifying its non-zero elements [Dempster, 1972]. Several research works have addressed this problem using penalization-based methods such as the graphical lasso [Yuan and Lin, 2007; Friedman et al., 2008] and corresponding Bayesian analogs [Armstrong et al., 2009; Baladandayuthapani et al., 2014; Zhang et al., 2016]. However, all of these methods make the canonical assumption of independence between the samples in estimating a (static) population-level precision matrix.

Over the last few years, several methods have been developed for dependent data via incorporation of sample-level information/covariates. Frequentist frameworks use penalty functions based approach to infer sample specific graphs and they are unable to quantify uncertainty [Liu et al., 2010; Zhang and Li, 2022]. Bayesian methods enable the estimation and inference of personalized networks but are often computationally intensive and limited in their ability to incorporate multiple covariates or capture nonlinear dependencies [Ni et al., 2019, Ni et al., 2022; Wang et al., 2022; Chen et al., 2025]. Under replicated settings, functional graphical modeling approaches have been proposed for analyzing multivariate functional data, which involves multiple measurements obtained across time points. Several research works have focused on dynamic graphical models with sliding window approaches [Kucyi and Davis, 2014], kernel-based nonparametric methods [Zhou et al., 2010; Gibberd and Nelson, 2017], and functional principal component framework [Zhu et al., 2016; Qiao et al., 2019; Zhang et al., 2021]. However, most of these methods are developed for univariate domain (i.e. time) and are not readily adaptable to spatial settings. Very few recent research works have been developed to study spatially varying correlation structures for multivariate spatial data [Lin et al., 2017; Dey et al., 2021] as well as ST data [Bernstein et al., 2022; Acharyya et al., 2022]. However, these methods are either limited to marginal associations or are unable to quantify spatially varying graphs (i.e precision matrices) over the entire spatial domain.

### Our contributions

In this paper, we propose a *spatial graphical regression* (`sGR`) model to infer spatially varying graphs for high-resolution multivariate spatial data. In contrast of existing methods, `sGR` can (I) explicitly incorporates spatial information to infer non-linear conditional dependencies; (II) enable flexible nonparametric estimation of spatial dependencies using Gaussian processes; (III) conduct estimation and global-local selection of spatially varying edges; (IV) scale with high dimensions, both in the number of spatial locations and features (genes). To induce parsimony, we employ a neighbourhood regression technique which provides a non-linear functional map from the spatial domain to a class of spatially varying precision matrices which engenders a spatially dependent conditional cross-covariance functions. We implement a scalable methodology through a conflation of spatially varying regression models and variable selection priors to induce sparsity. While we focus on undirected graphs, `sGR` can adapted to any general class of graphs (e.g. directed graphs under the known ordering of the nodes). `sGR` is motivated by and applied to two ST datasets from human breast and prostate cancer tissue to estimate spatially varying graphs, to identify spatial gene expression programs in the TME. To this end, we characterize a few key driver genes (genes with high degree of connectivity) for both cancer tissues; for example, we discover BIN2 as driver gene for the breast cancer tissue which prohibits tumor cell proliferation and multiple genes from ARHGAP family for the prostate tumor which regulates carcinogenesis. The general construction of `sGR` can be adapted to any class of directed and undirected spatial graphs and other scientific contexts.

### Sectional contents

We delineate the model construction of `sGR` in Section 2 followed by details of prior specification, posterior inference and hierarchical global-local inference in Section 3. The simulation study related results for undirected and real data based settings are provided in Section 4. Section 5 lays out the detailed application results and downstream analysis of our method on two ST datasets from breast and prostate cancer. Section 6 provides some conclusions and discussions. The Supplementary Materials contain additional details of methodology, simulation studies and ST data applications. A modular R package for fitting and visualization for `sGR` is available at https://github.com/******.

## 2 Spatial graphical models

Suppose the observed data is denoted as **Y**(**𝒮**) ^=^ [*<u>Y</u>*_1_(**𝒮**), …,*<u>Y</u>*_*p*_(**𝒮**)] (gene expression data in our context) which are observed over a spatial domain 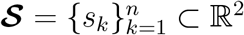 (tissue section in our context) – as shown in Figure 1A. For notational purposes, the matrices are denoted in bold and vectors are in underscore throughout the manuscript. Our inferential object of interest is a spatially varying graphical model, characterized by *G*(**𝒮**) = {*V, E*(**𝒮**)} where *V* is the vertex set of the *p* random variables and *E*(**𝒮**) denotes the spatially varying edge set which captures spatially varying conditional dependency among the variables. This engenders two primary inferential goals

(I) characterization of graph topology changing over spatial domain, *E*(**𝒮**), and

(II) identification of significant *global* (*E*(*s*) = 0, *∀ s ∈* **𝒮**) and *local* (*E*(*s*^*\**^) ≠ 0, where *s*^*^ ⊂ **𝒮**) spatial domains for spatially varying GGM.

We focus on undirected spatially varying Gaussian graphical model but it can be adapted to directed graphs with known order of nodes. To ease model exposition, we start with GGM in Section 2.1 which is generalized to a spatial graphical model and spatial graphical regression in Sections 2.2 and 2.3 respectively.

### 2.1 Gaussian graphical models

In a GGM, the presence (absence) of an edge between two variables indicates that they are conditionally dependent (independent) given all other variables. This conditional dependence is traditionally represented by the non-zero entries in the precision matrix associated with the GGM. Mathematically, a GGM is represented by an undirected graph *G* = (*V, E*) and the associated Gaussian distribution for p-dimensional random variable *<u>Y</u>* = (*y*_1_, …, *y*_*p*_) is *N*(0, **Ω**^*−*1^) where the precision matrix **Ω** = (*ω*_*ij*_)_*p* × *p*_ encodes the conditional dependence between variables. Particularly, *ρ*_*ij*_ = Corr^(^*<u>Y</u>*_*i*_, *<u>Y</u>*_*j*_ | *<u>Y</u>*_*l*_, *l* ≠ *i, j*) denotes the partial correlation between*<u>Y</u>*_*i*_ and *<u>Y</u>*_*j*_ given all other variables. The nodes {*i, j*} are connected to each other iff *ρ*_*ij*_ ≠ 0. The partial correlation can be expressed in terms of elements of the precision matrix as 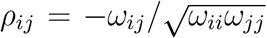. This implies that *ρ*_*ij*_ ≠ 0 (= 0) is equivalent to *ω*_*ij*_ ≠ 0 (= 0) where *ω*_*ij*_ is the (*i, j*)th element of the precision matrix **Ω**. The *ω*_*ij*_ ≠ 0 (= 0) leads to conditional dependence (independence) between *i*-th and *j*-th node. The undirected graph structure can now be represented as *G* = (*V, E*) with vertex set *V* = {1, …, *p*} and edge set *E* = {(*i, j*) : *ρ*_*ij*_ ≠ 0, *i* ≠ *j*} = {(*i, j*) : *ω*_*ij*_ ≠ 0, *i* ≠ *j*}. Therefore, estimation of graph structure boils down to precisely identifying the non-zero locations of the precision matrix based n observed realization of p-dimensional vector*<u>Y</u>* .

### 2.2 Spatially varying Gaussian graphical models

In this paper, we consider a spatially varying Gaussian graphical model. For a given spatial location *s* (*∈* **𝒮** ⊂ ℝ^2^), we assume that the joint distribution of *<u>Y</u>* (*s*) = [*y*_1_(*s*), …, *y*_*p*_(*s*)] is a multivariate normal distribution

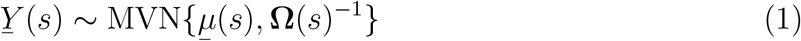

where **Ω**(*s*) = {*ω*_*ij*_ (*s*)} is a *p* × *p* spatially varying precision matrix and our primary estimand of interest. Analogously to GGM, consider the partial correlation between*<u>Y</u>*_*i*_(*s*) and *<u>Y</u>*_*j*_ (*s*) is *ρ*_*ij*_ (*s*) = corr(*<u>Y</u>*_*i*_ (*s*), *<u>Y</u>*_*j*_ (*s*) |*<u>Y</u>*_*l*_(*s*), *l ∉* {*i, j*}). The spatially varying graphical model is formulated as,

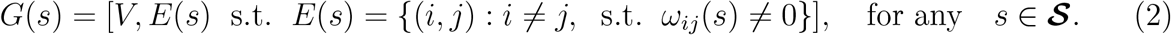

Under the multivariate normal assumption, for any *s ∈* **𝒮**, *ω*_*ij*_ (*s*) = 0 is interpreted as the spatially varying nodes *<u>Y</u>*_*i*_(*s*) and *<u>Y</u>*_*j*_ (*s*) are conditionally independent given all other nodes *<u>Y</u>*_*l*_(*s*) at location *s* i.e*<u>Y</u>*_*i*_(*s*) *⫫ Y*_*j*_ (*s*) |*Y*_*l*_(*s*) s.t. *l* ≠ {*i, j*}. Our goal of this paper is to characterize the spatially varying partial correlation function *ρ*_*ij*_ (*s*) for a specific location *s ∈* **𝒮** with elements spatially varying precision matrix such as *ω*_*ij*_ (*s*) and *ω*_*ii*_(*s*). Note that *G*(*s*) is defined for each location *s* and represents a spatially-varying graph over the whole space **𝒮**. In equation (1), we specify that *<u>Y</u>* (*s*) follows a multivariate Gaussian distribution and it is worth noting that *<u>Y</u>* (*s*) and *<u>Y</u>* (*s*^*′*^) can be correlated for any two locations {*s, s*^*′*^} *∈* **𝒮**.

There are two <u>desiderata</u> for spatial graphical models: 1) enable parsimony or sparsity in the graphical structure – to allow tractability in estimation, computations and interpretability of the edges; and 2) to admit a flexible structure on *ω*_*•*_(*s*) (or equivalently *ρ*_*•*_(*s*)) to capture various types of spatial dependencies (global and local). We achieve these two goals by casting this model in spatial graphical regression framework as discussed next.

### 2.3 Spatial graphical regression model

The spatially varying precision matrix **Ω**(**𝒮**) = {**Ω**(*s*_1_), …, **Ω**(*s*_*n*_)} is a high-dimensional object consisting of *n* × *p* × (*p −* 1)*/*2 edges (*≈* 2 × 10^6^following dimensions from our real data applications) which makes the joint estimation challenging and may even make it infeasible with high dimensions of *n* and *p*. To address such ultra high-dimensional problem, we develop a spatially varying neighborhood selection framework to infer the non-zero locations of the spatially varying precision matrix **Ω**(**𝒮**) which encodes the spatial varying conditional structure of the graphs. The neighborhood selection framework has been used to estimate covariate-dependent precision matrices in context of non-spatial graphical models [Meinshausen and Bühlmann, 2006; Ni et al., 2019; Ha et al., 2021; Zhang and Li, 2022; Wang et al., 2022]. Following the sparse nature of the spatial genomics data, the effective number of edges can be reduced by incorporating variable selection technique and flexibility of the framework is ensured through non-parametric modeling structure as discussed next.

To this end, we introduce a novel spatial graphical regression (`sGR`) method to infer spatially varying neighborhoods for an undirected spatial graphical model. `sGR` is developed using Gaussian processes and provides a non-linear functional map between spatially dependent non-parametric regression coefficients and spatially varying precision matrices.

For the rest of the paper, we are going to assume *<u>µ</u>*(*s*) = 0 *∀s ∈* **𝒮** since we can scale the data and set the mean to be zero. We denote the i-th node as *<u>Y</u>*_*i*_(**𝒮**) = (*y*_*i*_(*s*_1_), …, *y*_*i*_(*s*_*n*_)) *⊤* and provide the node-specific undirected spatial graphical regression on 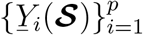 as

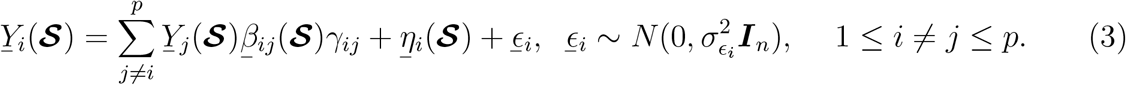

In the model above,*<u>β</u>*_*ij*_ (**𝒮**) is the spatially varying regression co-efficient between (*i, j*)-th gene and *γ*_*ij*_ is the selection indicator which regulates the existence of spatially varying edge on the whole spatial surface. The parameter *<u>η</u>*_*i*_(**𝒮**) captures the spatial proximity based dependency for *i*-th gene and ϵ_*i*_ denotes heteroscedastic errors which is modeled with a Gaussian distribution with standard error 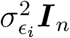.

#### **Characterization of** *<u>β</u>*_*ij*_ (**𝒮**)

We define *<u>β</u>*_*ij*_ (*s*) as the <u>spatial conditional precision function (SCPF)</u> for a specific location *s ∈* **𝒮**. The event *I*{*<u>β</u>*_*ij*_ (**𝒮**) *≡* 0} determines the spatially varying graphical structure and encodes the conditional independence between nodes {*<u>Y</u>*_*i*_(**𝒮**),*<u>Y</u>*_*j*_ (**𝒮**)}_(1*≤i*≠*j≤p*)_ over the spatial domain **𝒮**. Recall that the spatially varying partial correlation function between two nodes *<u>Y</u>*_*i*_(*s*) and *<u>Y</u>*_*j*_ (*s*) for a specific location *s* is quantified with *ρ*_*ij*_ (*s*) (1 *≤ i* ≠ *j ≤ p*) where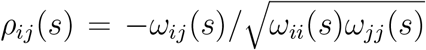. The edge selection rule then suggests that an edge (*i, j*) *∈ E*(**𝒮**) (as defined in equation (2)) is significant if and only if *ρ*_*ij*_ (*s*) ≠ 0 for any *s ∈* **𝒮**.

In the context of GGMs, Peng et al. [2009] showed that the partial correlation between two nodes can be characterized by their corresponding regression coefficient (Lemma 1 in their paper). We propose a spatial analog of their lemma in the context of spatial graphical regression framework, which can stated formally as:

**Lemma 1**: *Under the regression framework of node y*_*i*_(*s*) *on other nodes y*_*−i*_(*s*) *for a specific location s ∈* **𝒮**, *the spatially varying-coefficient model is* 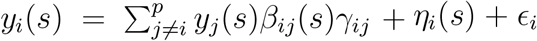. *The underlying assumption: η*_*i*_(*s*) + _*i*_ *is uncorrelated with y*_*−i*_(*s*), *if and only if* 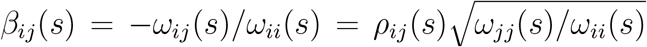 *for any s ∈* **𝒮** *where ω*_*ij*_ (*s*) *and ω*_*ii*_(*s*) *denote off-diagonal and diagonal elements of* **Ω**(*s*) *respectively. Therefore the spatially varying partial correlation function can be expressed as* 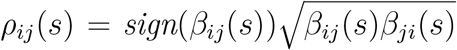 *and var*(*η*_*i*_(*s*) + _*i*_) = 1*/ω*_*ii*_(*s*) *for any s ∈* **𝒮**.

Lemma 1 suggests that the partial correlation *ρ*_*ij*_ (*s*) varies spatially as well as the regression coefficient of *y*_*i*_(*s*) over *y*_*j*_ (*s*) based on the spatial function *β*_*ij*_ (*s*) (*s ∈* **𝒮**). The SCPF *β*_*ij*_ (*s*) defines the relationship between the partial correlation and spatial locations through *ω*_*ij*_ (*s*) and *ω*_*ii*_(*s*) which are elements of the spatially varying precision matrix **Ω**(*s*). In this sense, the SCPF can be characterized as *β*_*ij*_ (*s*) = *−ω*_*ij*_ (*s*)*/ω*_*ii*_(*s*) which encodes the spatially varying conditional dependency structure of undirected graph for a specific location *s ∈* **𝒮**. The spatial random effect *η*_*i*_(.) captures spatial dependent structure across locations for particular node and characterizes the variance for *i*th node over the spatial domain **𝒮**. In essence, *β*_*ij*_ (*s*) and *ω*_*ii*_(*s*) = 1*/*var(*η*_*i*_(*s*) + _*i*_) characterizes the spatially dependent partial correlation function as defined in (1).

#### **Flexible structure for** *<u>β</u>*_*ij*_ (.) and *<u>η</u>*_*i*_(.)

As mentioned before, one of the desiderata is to have flexible model on the spatially varying edges – to capture the varying degrees of spatial dependency. Another challenge is that we are dealing with a spatially varying neighborhood selection problem (of order *O*(*np*^2^)). To enable such adaptable structure, we model *<u>β</u>*_*ij*_ (.) and *<u>η</u>*_*i*_(.) with spatial (kernel-based) Gaussian processes defined as,

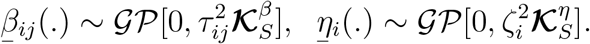

Spatially dependent Gaussian processes are standard technique in case of developing flexible methods through non-parametric spatial functions [Gelfand and Schliep, 2016; Reich and Fuentes, 2015].

#### Imposing sparsity through *γ*_*•*_

The underlying biological system of genomic networks is expected to sparse and leptokurtic with small number of large and spatially varying signals while most of the regression coefficients are small or effectively zero [Acharyya et al., 2022; Bernstein et al., 2022]. To adapt for sparse structure of the ST data, we transform the spatial graphical regression problem to a variable (i.e. edge) selection problem in a spatial context. To this end, we introduce the selection indicators *γ*_*•*_ which controls the sparsity of the spatially varying graphs and *γ*_*•*_ does not depend on spatial structure. More precisely, *I*(*γ*_*•*_ = 1) = *I*(*<u>β</u>*_*•*_(*s*) ≠ 0 ∃ *s* ∈ **𝒮**). This variable selection strategy, that has been previously employed for standard regression models (Kuo and Mallick [1998]; O’Hara and Sillanpää [2009]), helps us achieve three major goals:(I) straightforward estimation and inference on significant and insignificant spatially varying edges, (II) development of computationally efficient model which reduces cost of prediction and (III) increase in precision in case of parameter estimation [Mitchell and Beauchamp, 1988]. This formulation helps us to build a parsimonious and computationally efficient method with flexible priors to precisely estimate significant spatially varying edges (Section 3) followed by global and local inference on the spatially varying edges (Section 3.2), as we discuss next.

## 3 Bayesian inference

To complete the spatial graphical model specification, we discuss the priors of two primary components of the model in (3). First, we model the SCPF *<u>β</u>*_*ij*_ (.) using a Gaussian processes with spatial kernel and analogous steps are followed to model the spatial random effect *<u>η</u>*_*i*_(.) and second, we detail the variable selection variable to incorporate sparse data structure.

### 3.1 Gaussian process formulation of SPCF

Utilizing Mercer’s theorem [Williams and Rasmussen, 2006], we use Karhunen-Loéve expansion on *<u>β</u>*_*ij*_ (**𝒮**) and *<u>η</u>*_*i*_(**𝒮**) to decompose the Gaussian processes following Shi and Kang [2015] as follows:

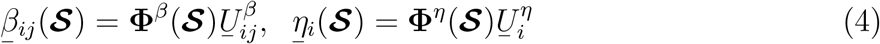

where **Φ**^*β*^(**𝒮**) is a *n* × *L*_*β*_ dimensional matrix containing basis vector and 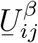 is a vector of length *L*_*β*_. The approximate number of basis coefficients is denoted with *L*_*β*_. Shi and Kang [2015] showed that under some mild regularity conditions, one can decompose the spatial kernel as 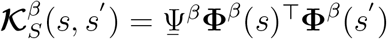 where <u>Ψ</u>^*β*^ denotes the eigenvalues of the kernel.

The orthogonality is ensured through the assumption based on the following conditions 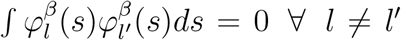 where 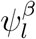 and 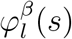 are elements of <u>Ψ</u>^*β*^ and **Φ**^*β*^(*s*) respectively. We use an analogous decomposition of the Gaussian process of *<u>η</u>*_*i*_(**𝒮**). Based on decomposition on equation (4), we place independent Gaussian priors on the basis coefficients 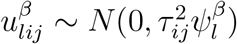 such that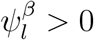. In spatial context, we employ modified squared exponential kernel [Shi and Kang, 2015; Wu et al., 2024; Lin et al., 2023] expressed as,

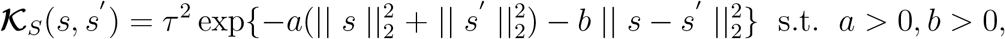

Where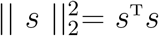. This implies 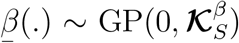 with following properties 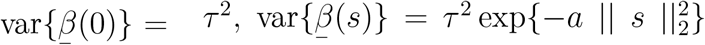 and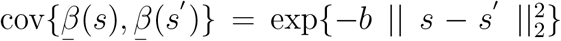. The scalar *τ*^2^controls the marginal variance which is maximum at origin 0. The parameter *a* controls the decay rate var{*<u>β</u>*(*s*)} in comparison with var{*<u>β</u>*(0)} which means large values of *a* implies high decay rate. The parameter *b* modulates the smoothness of the function *<u>β</u>*(*s*). Without loss of generality, one can set *τ*^2^ = 1 and the MSE kernel is simplified to a squared exponential kernel by setting *a* = 0. Further details about the spatial MSE kernel decomposition and the construction of the basis vectors is deferred to the Section S1.1 of the Supplementary Materials.

#### Decoupling of *<u>β</u>* and *γ*

We consider the independent prior structure and posit a Bernoulli prior on *γ*_*ij*_ *∼* Ber(0.5) along with a Gaussian process prior on SCPF *<u>β</u>*(.) as discussed in the previous paragraphs. The independent prior structure has several advantages over the dependent structure. (I) The sampling algorithm to fit the model does not require any tuning parameter because of the independent priors on (*<u>β</u>, γ*) A practical challenge while implementing variable selection through spike-and-slab prior is finding the right model tuning, which involves adjusting for the prior distribution to ensure that the Markov chain Monte Carlo (MCMC) chains mix well, enabling the sampler to efficiently transition between the slab and spike. (II) The independent priors also help with computational efficiency. In case of spike-and-slab prior, one needs to marginalize *<u>β</u>* to obtain the conditional update of the selection variable *γ*. This process results to inversion *n* × *n* kernel matrix for every iteration, which is significant computationally intensive. (III) One can incorporate prior information through variable selection technique which will encourage genomics association across multiple spatial regions. The independent prior structure [Kuo and Mallick, 1998] has found several applications in genetics [Sillanpää and Bhattacharjee, 2005].

To complete the prior specification parameters, we place conjugate prior on error variance 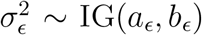 where IG(*a, b*) represents an inverse gamma distribution with shape a and rate b. In the next Section 3.2, we layout the posterior sampling algorithm and global-local inference of the SCPF for a significant spatially varying edge i.e. *I*{*γ*_*ij*_ = 1}.

### 3.2 Posterior inference

#### Sampling algorithm

For full Bayesian inference on model (3), we develop an efficient MCMC algorithm for posterior inference of the following parameters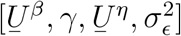. The algorithm is a combination of Gibbs sampler 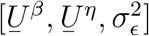 and an accept-reject algorithm [*γ*]. For computational efficiency, we fit the spatial graphical model with node wise parallel regressions and the sampling algorithm doesn’t necessitate any tuning parameter. However, in case of *I*{*γ*_*ij*_ = 0}, the updated value of SCPF is sampled from the full conditional distribution which which effectively simplifies to its prior distribution. We draw the samples of SCPF from it’s posterior distribution when *I*{*γ*_*ij*_ = 1} s.t. 1 *≤ i* ≠ *j ≤ p*. We use the accept-reject algorithm to update the value of the selection indicator. Additional details of the sampling algorithm can be found in the Section S1.2 and a graphical representation of the `sGR` model is provided in Figure S1 of the Supplementary Materials.

#### Positive definiteness property

The estimated regression coefficients from the spatial graphical regression methodology (3) does not ensure the symmetry and positive definiteness property of the spatial precision matrices (**Ω**(**𝒮**)^*−*1^). We employ the well-established OR rule to enforce symmetry of the precision matrices in standard (non-spatial) graphical model settings [Meinshausen and Bühlmann, 2006]. More specifically, {*i, j*}-th node is included if one of these three events occur *I*{*γ*_*ij*_ = 1, *γ*_*ji*_ = 1}, *I*{*γ*_*ij*_ = 1, *γ*_*ji*_ = 0}, and *I*{*γ*_*ij*_ = 0, *γ*_*ji*_ = 1} (excluded when *I*{*γ*_*ij*_ = 0, *γ*_*ji*_ = 0}). We consider the estimate the SCPF by selecting the higher value of posterior inclusion probability between *I*{*γ*_*ij*_ = 1} and *I*{*γ*_*ji*_ = 1} which works well in practice to study genomics association [Wang et al., 2022; Zhang and Li, 2022].

To ensure positive definiteness property, we need the **Ω**(*s*) to be a diagonally dominant matrix *∀s ∈* **𝒮**. We implement a rescaling step in post processing procedure of posterior samples to ensure positive definiteness of the estimates of spatially varying precision matrices [Cheng et al., 2014; Zhang and Li, 2024]. We denote 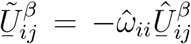 where 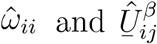 are estimated values of varaince and basis coefficients from (3) and (4). We set 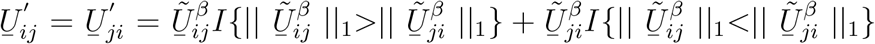 and enforce symmetry of the basis coefficients which will automatically ensure the symmetry of the SCPFs. Next, we focus on the rescaling step to maintain the positive definiteness property. We assume that the true precision matrix is diagonal dominant and **Φ**^*β*^(**𝒮**) is within known ranges i.e. || **Φ**^*β*^(**𝒮**) ||*∞*= *a < ∞*. We rescale and set the final estimate of 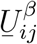 to 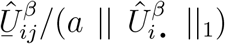 where 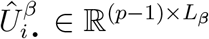 is a matrix containing basis coefficients. This rescaling step ensures diagonal dominance of the final estimator along with the positive definite property and does not impact the sparse structure. Step-by-step derivations of this rescaling step is provided in the Section S1.3 of the Supplementary materials. Next, we perform a global-local inference procedure on the selected spatially varying edges and identification of the hub nodes (genes) with high connectivity degree as shown in Figure 1C.

#### Hierarchical global and local inference

We develop a hierarchical (two level) global and local inference to infer and interpret the spatially varying edges using multiple testing corrections. For the first level selection, our hypothesis is two nodes (i,j) are spatially independent over the <u>entire spatial domain</u> based on the event *I*{*γ*_*ij*_ = 0} and we define the global edge probability (GEP) as

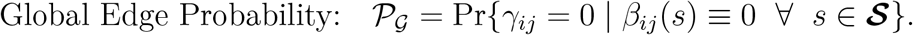

The event *I*{*γij* = 0} indicates *β*_*ij*_ (**𝒮**) ≡ 0 which suggests that the SCPF is 0, implying that there are no significant edge between two nodes (*i, j*) over the whole spatial domain **𝒮**. In case of the value of GEP *<* 0.5, we focus on the counter event *I*{*γ*_*ij*_ = 1} which implies the spatially varying edge between two nodes is significant over some compact subset of the spatial domain, which encompasses the global selection of a spatially varying edge. Once these “global” edges are determined, subsequently, we aim to identify the local sub-regions of the spatial domain where the spatially varying edge is significant (jointly) controlling for family-wise error rate.

To this end, we introduce the local edge probability (LEP) in case of {*γ*_*ij*_ = 1} which indicates *β*_*ij*_ (*s*) ≠ 0 for some *s* ∈ **𝒮**. This feature enables us to identify the local spatial sub-regions where the spatial curve is bounded away from zero and controls for multiplicity correction factors. Specifically, following steps from Ruppert et al. [2003], we construct 100(1 − *α*)% joint credible bands for *β*_•_(*s*) (*j* ≠ *i* | *i, j* ∈ {1, …, *p*}) as

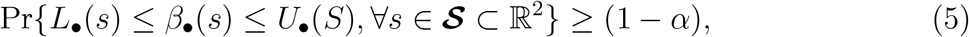

where *L*_•_(*s*) and *U*_•_(*s*) the lower and upper bound respectively for the undirected setting while i-th node is being regressed on j-th node (*j* ≠ *i*). An interval satisfying (5) can be expressed as 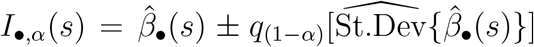, where 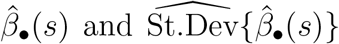 are the mean and standard deviation for a given *S* taken over all the MCMC samples.

Finally, we invert these joint credible bands to construct *I*_•,*α*_(*s*) for multiple levels of *α* and determine for each *s* the minimum *α* at which each interval excludes zero, denoted *P*_•,LEP_(*s*) = min{*α* : 0 ∉ *I*_•,*α*_(*s*)}, which can be computed by

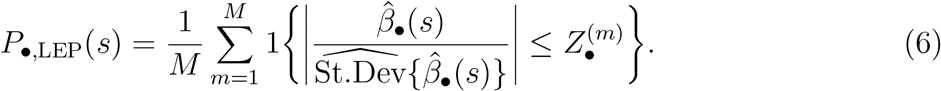

The *Z* is constructed in spatial context following Meyer et al. [2015] and additional details 16 are provided in Section S1.4 of the Supplementary materials. These local probability scores have global properties, particularly to adjust for multiple testing i.e. for a specific *α*, we identify *s* for which *P*_•,LEP_(*s*) *< α* as significant. This is equivalent with checking the zero inclusion or exclusion of the joint credible interval for a specific value of *α*.

## 4 Simulation studies

To showcase the effectiveness of `sGR`, we perform replicated simulations across different settings to assess the performance in terms of graph structural recovery. We first present the data generation mechanism and the comparative methods followed by the details of evaluation on graph recovery rates for both undirected (Case I) and real data based simulation settings (Case II) below. All the results are summarized over 50 replicates. We run 5000 iterations and discard the first 50% of the posteriors sample obtained from the sampling algorithm and posterior inferential techniques are detailed in Section 3.

### Data generation scheme

For a given spatial location *s* (∈ **𝒮** ⊂ ℝ^2^), we generate the data *<u>Y</u>* (*s*) = (*y*_1_(*s*), …, *y*_*p*_(*s*)) from a multivariate normal distribution with spatially varying precision matrix. Without loss of generality, the spatial locations are generated from a square *s* ∈ [−1, 1]^2^ and data *<u>Y</u>* (*s*) ∼ *N*_*p*_{0, **Ω**(*s*)^−1^} where **Ω**(*s*) denotes the true precision matrix at the spatial location *s*. Following the real data dimensions, we set (*n, p*) = (5000, 100) for the <u>undirected spatial graph settings</u> and generate the data from the undirected `sGR` model (3) with 2% significant spatially varying edges. We define the signal-to-noise ratio (SNR) for the spatial graphical regression w.r.t. specific node *i* as 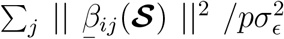. For all the simulations in the manuscript, we set SNR=3. The values of node specific noise variances are generated from uniform(0.2,0.4). For the <u>real data bases simulation settings</u>, we vary number of spatial locations *n* = {3000, 4000, 5000, 6000, 7000} and set number of genes/features *p* = 100 with 2% significant edges. For data generation purposes, we simulate *<u>Y</u>*_1_(**𝒮**) ∼ *N*(0, 1) and 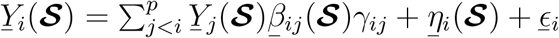 where 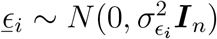 and *i* = 1, …, *p*. We consider different spatial functional forms for SCPF which are listed in Section S2 of the Supplementary Materials and the spatial pattern is being varied across the spatial domain of the tissue.

Comparative methods: To the best of our knowledge, there are no existing methods for direct comparative assessment. However, we compare our proposed methodology with the following relevant graphical-modeling approaches adapted for the spatial setting in the following manner. (I) GraphR [Chen et al., 2025] is a probabilistic graphical regression method with linear external covariates and we treat the spatial co-ordinates as linear covariates for our comparison study. (II) Zhang and Li [2022] proposed a class of penalized graphical regression methods in high-dimensional setting with linear covariates. We set the spatial co-ordinates to be linear covariates to use as a frequentist penalization-based method to compare with our `sGR` method. (III) A non-linear method, Graph-optimized classification and regression trees (Go-CART, Liu et al. [2010]) which is a covariate dependent semi-parametric regression-based model to estimate undirected graphs. (IV) An adaptation of the standard Graphical Lasso (Glasso, [Friedman et al., 2008]) to incorporate spatial information, as follows: we implement Glasso in a localized spatial structure with a fixed radii (of 0.25) and the structural recovery measures are calculated aggregating across all such neighbourhoods. The reported measures are averaged for all the neighbourhoods over the whole spatial domain.

### Evaluation metrics

We use the following comparative measures to evaluate the graph structural recovery performance across different methods such as true positive rate (TPR), false positive rate (FPR), false discovery rate (FDR) and Matthews correlation coefficient (MCC). The definition of structural recovery rates are deferred to Section S2 of the Supplementary Materials. MCC [Matthews, 1975] is a measure of binary classification where values ranging from −1 (total disagreement) and +1 (perfect classification). We also construct the receiver operating characteristic (ROC) curve to compare the compare the performance of different methods in terms spatially varying structural recovery. The area under the ROC curve (AUC) is computed by calculating sensitivity (TPR) and 1-specificity (FPR) at each threshold parameter value in between [0, 1] and used as a comparative measure across different methods. High values of AUC and MCC implies better structural recovery of spatially varying edges.

### Case I: Undirected spatial graphical models

In this section, we present the results from detailed simulation study to assess the performance of the spatial graphical regression method in case of spatially varying graph structural recovery in undirected setting. Figure 2 shows the performance of undirected spatial graphical regression method in terms of graph structural recovery from comparative study after averaging over 50 replicates. The `sGR` method outperforms the competitive methods in terms of AUC (`sGR`: 0.97, GraphR: 0.87, Go CART: 0.85, Penalized GR: 0.73, Glasso: 0.69) and MCC (`sGR`: 0.93, GraphR: 0.79, Go CART: 0.76, Penalized GR: 0.62, Glasso: 0.58). The values of other evaluation matrices of graph structural recovery (TPR, FDR, FPR) is reported to Section S2 of the Supplementary Materials. The `sGR` method has obtained the highest values of AUC, MCC, and TPR with the lowest values of FDR and FPR while comparing with other methods. Rest of the methods has high values of TPR and comparatively high values of FPR as well which results into a relatively low value for MCC. This relationship can be explained as trade-off between TPR and FDR. The GraphR method is based neighbourhood based regression technique but it can only incorporate spatially varying linear dependence. Go CART and Penalized GR are optimization and penalized regression based frequentist methods which are unable to do uncertainty quantification. Glasso was implemented based spatial sub-regions which is unable to recover the underlying spatially varying graph structure. The `sGR` method achieves better results in terms of graph structural recovery compared to the competing methods in case of spatially varying graph scenarios.

**Figure 2.**
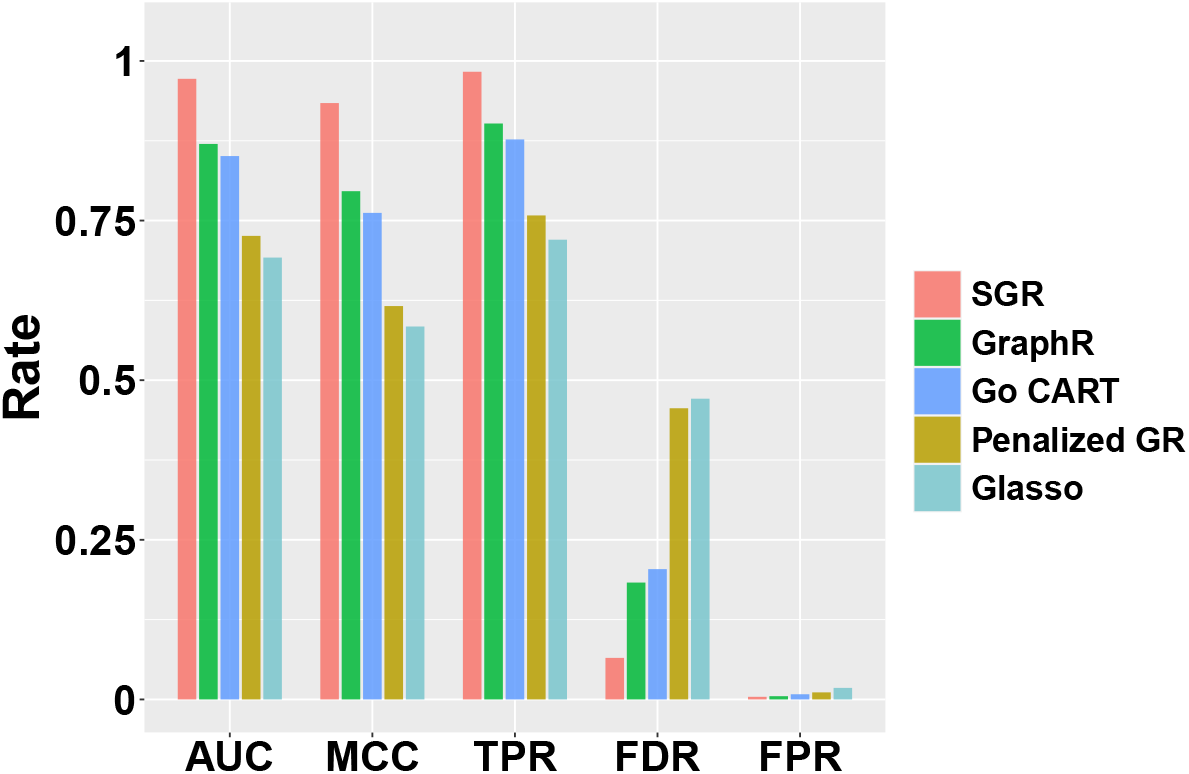
Comparative study of sGR in undirected settings: To assess the selection performance of the undirected spatial graphical models, we conduct a simulation study across 50 replicates and report the mean values for evaluation metrics such as AUC, MCC, TPR, FPR, and FDR for every undirected setting with (*n, p*) = (5000, 100) and SNR= 3.

### Case II: Real data based simulation settings

First, we assess the performance of `sGR` model in case of real data based simulation settings with known order of nodes w.r.t. structural recovery and estimation along with global local inference. Mimicking the real data dimensions, we consider (*n, p*) = (5000, 100) to analyze the estimation and selection performance. Our underlying hypothesis is that the `sGR` method is able to precisely estimate and recover different spatial patterns. Figure 3A shows the performance of our method in terms of graph structural recovery for the directed setting. The right panel of Figure 3A illustrates the true network consisting of 12 (non-zero) spatially varying edges and an additional 12 non-existent edges. The subsequent panel depicts each corresponding edge’s GEP, with the edge widths being directly proportional to the GEP. The final panel indicates the binary selection of the estimated edges, which is dependent on their individual GEP, and compares it with the standard threshold 0.5 as discussed in Section 3. Figure 3A indicates that the directed spatial graphical model is able to recover all spatially varying edges i.e. SCPFs. Next, we focus on the estimation and global-local inference of SCPFs.

**Figure 3.**
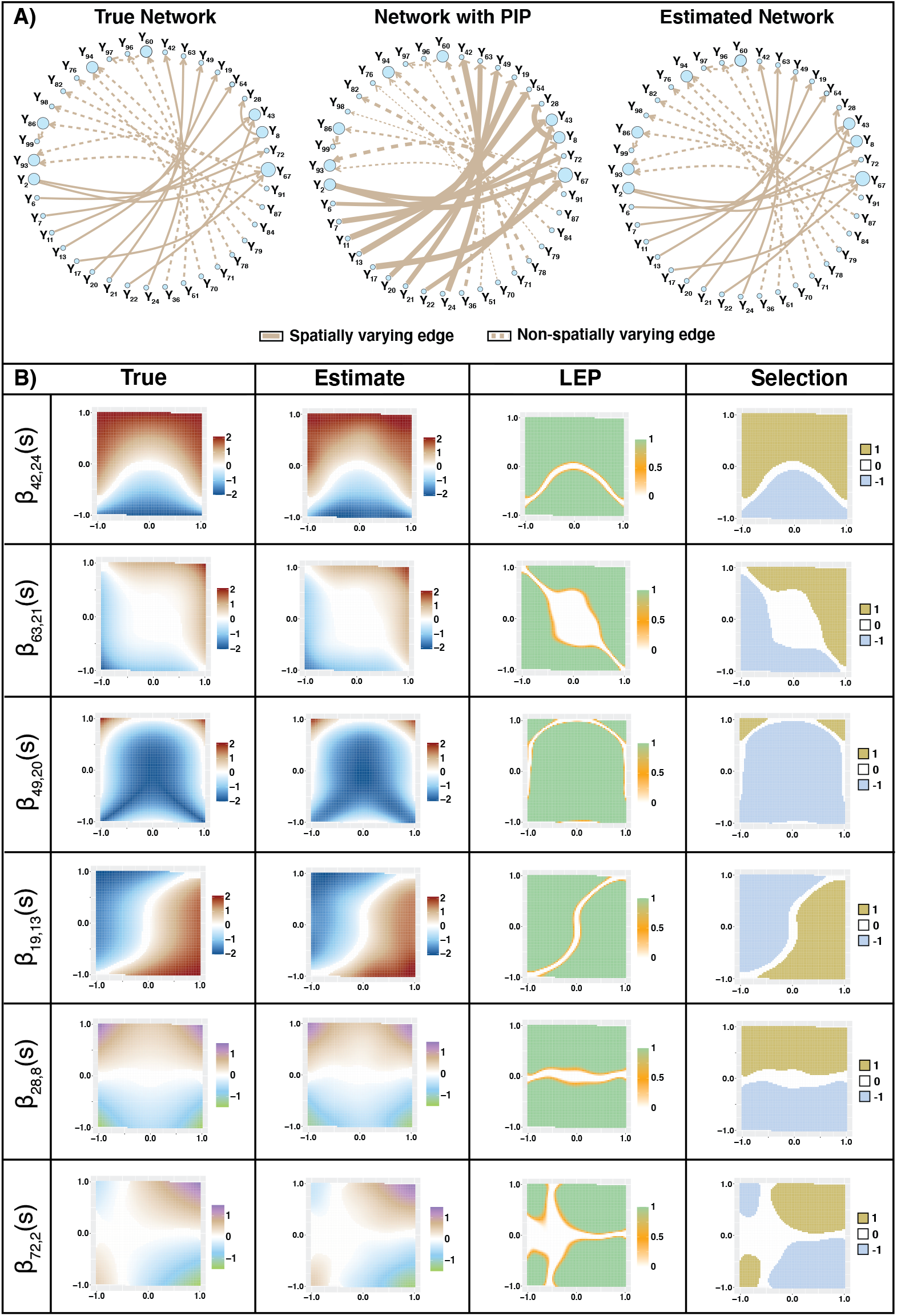
Performance of sGR under real data based settings: The left graph in Panel A) shows the true network with 12 spatially varying edges and randomly chosen another 12 non-existent edges. The middle graph represents GEP of each edge where edge-widths are proportional to the GEP. The right graph is the binary selection of the estimated edges based on their respective GEP. Panel B) shows 6 spatially varying edges (*β*_42,24_, *β*_63,21_, *β*_49,20_, *β*_19,13_, *β*_28,8_, *β*_72,2_) out of 12 as mentioned in Panel A). The first, second, third and fourth column of the plot represents true, estimate, LEP and selection of the SCPFs respectively.

Figure 3B shows the estimation performance of the `sGR` model under known order of nodes for 6 SCPFs with different spatial patterns. The spatial patterns are considered according to the morphological characteristics of different tissue based on real data. The functional forms are mentioned in Section S2 of the Supplementary Materials. The first and second column of Figure 3B contains the true and estimated SCPFs before applying SimBaS method as discussed in Section 3.2. In the third column of the figure we have the spatially varying LEP for each spatial location. More specifically, the third column displays the *P*_•,LEP_ from equation (6) across the spatial domain which depicts the spatial regions where SCPFs are bounded away from 0. The final column shows the corresponding spatially varying local selection and sign of SCPFs over the spatial domain. The `sGR` model can recover the graph structure and jointly estimate different SCPFs under the real data based simulation settings. Analogous results for the rest of the 6 SCPFs are provided in Figure S2 of the Supplementary Materials.

### Additional simulations

We conducted additional simulations across multiple SNRs (ranging from SNR = {1, 3, 5}), results of which are provided in Section S2 of the Supplementary Materials. We assess the consistency performance of `sGR` in terms for estimation and graph structural recovery with varying spatial locations and a fix number of nodes in Figure S15 of the Supplementary Materials. The `sGR` model with known order of nodes performs better in terms of graph estimation and structural recovery with increasing number of spatial locations.

## 5 Network estimation of spatial transcriptomics data

Our `sGR` method is motivated by and applied to two ST datasets from human breast (Section 5.1) and prostate cancer tissues (Section 5.2). These datasets are collected through the 10X Genomics Visium platform. In addition to spatial gene expression data, both tissue sections are annotated with tumor, intermediate, and normal regions. This annotations are readily accessible on the 10X Genomics website which we use solely for interpretational purposes. Additional details of data collation and preprocessing are provided in Section S3.1 of the Supplementary Materials.

### Scientific goals

Our scientific goal here is to identify global and local region based spatial connectivity between genes in the TME. Accurate spatial delineation of these gene expression connectivity can shed light on conserved and differential co-regulation patterns in different areas of the tumor tissue (e.g. tumor vs. normal). This can provide deeper insights into the molecular mechanisms driving cancer evolution, identification of driver genes and guiding the development of targeted therapies, especially immuno-oncological drugs. To this end, we focus our analyses on subset of immune signature genes linked to immune cell populations in human tissues Nirmal et al. [2018]. For both breast and prostate cancer datasets, we specifically investigate the connectivity of B-cell and T-cell immune signature genes and their associated functional properties in the TME.

B-cells generate antibodies to fight infections, while T-cells protect individuals by eliminating cancerous and infected cells, preventing infections [Tan et al., 2022]. Understanding the functionality of immune signature genes related to B-cell and T-cell in a tumor environment is crucial for developing immunotherapies, a rapidly growing sector in cancer treatment. By boosting the body’s natural defenses (B-cells’ antibody production and T-cells’ destructive capabilities), immunotherapy can drive the path for innovative cancer management strategies, and paves the way for personalized cancer therapy.

### 5.1 Breast cancer

The human breast cancer tissue sample: an estrogen receptor-positive, progesterone receptor-negative, human epidermal growth factor receptor (HER)2-amplified (HER+) invasive ductal carcinoma was prepared on the Visium platform with immunofluorescence staining. The immunofluorescent image of the breast cancer tissue section, with *n* = 4, 727 spots, is showed in Figure 4A. Pathologist annotated the invasive carcinoma tissue sample as tumor, intermediate and normal region. The regions delineated by red and blue boundaries are classified as tumor and normal areas respectively, with the remaining tissue area being marked as intermediate [Zhao et al., 2021].

**Figure 4.**
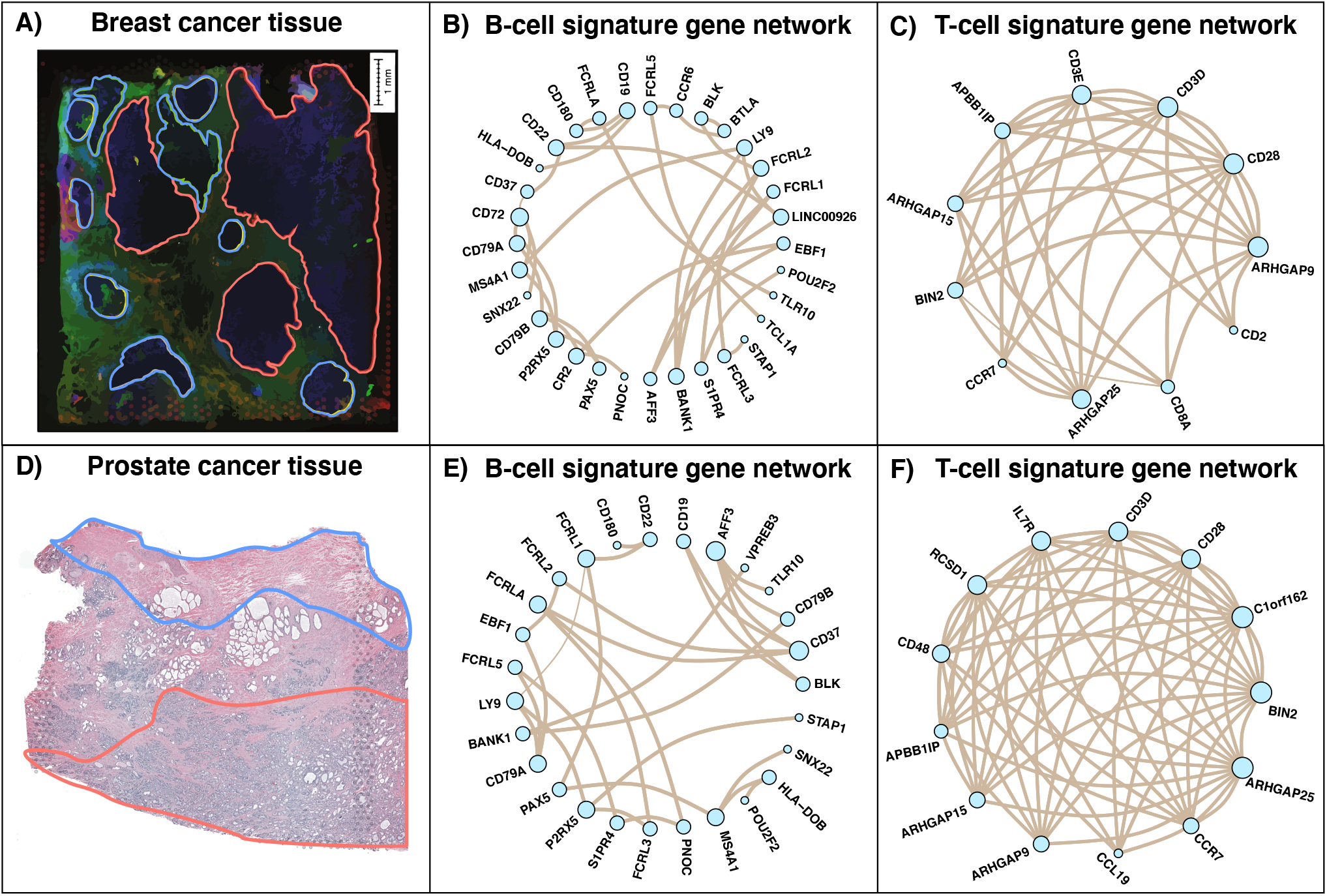
Spatial distribution and global network of breast and prostate cancer tissue: Panel A) shows the immunofluorescent image of the breast cancer tissue with 4,727 spots and the corresponding histopathological annotations. Panel B) shows the hematoxylin and eosin staining image of the human prostate cancer tissue with 4371 spots. The red and blue colored regions denote the tumor and normal regions of the tissue respectively. The remaining area within the tissue is regarded as transitional or intermediate phase. The network of immune signature genes for (B, T) cells across the entire tissue is displayed in Panels B) and C) for breast and Panels E) and F) for prostate cancer tissue respectively.

In the breast cancer data, we found 33 and 79 immune signature genes associated with B-cell and T-cell respectively following Nirmal et al. [2018]. We apply the `sGR` method (Section 2.3) on B-cell and T-cell signature genes to obtain the networks on the whole tis-23 sue domain as displayed in Figures 4B and 4C respectively. To parse out the dense network, we only show the network of T-cell signature genes with high values of scaled connectivity degree (*>* 0.5) in Figure 4C. We define the scaled connectivity degree as number of connected edges for each gene and scale the values with the maximum value. The network plot considering all the genes is provided in Figure S16 of the Supplementary Materials. These networks show the global connectivity among the B-cell and T-cell signature genes and obtained by following the posterior inference steps as discussed in Section 3.

We further dissect the networks and observe some of the spatially varying edges related to B-cell and T cell-signature genes in top and bottom row of Figure 5 respectively. To assess connectivity across tissue regions, we compute the scaled adjusted rand index (sARI is defined in Section S3 of the Supplementary Materials) between the positive/negative interaction of each edge and the histopathological annotation (tumor, intermediate or normal) of the spatial regions. We report the edges with high values of sARI and overlay the spatial region annotations for the LEP (*P*_•,LEP_(*s*)) and positive/negative interaction plot. The sARI-based rank for spatially varying edges helps us with our scientific goal to identify the change in gene-gene local spatial connectivity in TME. Figure 5, the top row contains the estimated spatially varying edge, LEP and selection for FCRL2 − BLK, TCL1A − FCRL5, CD180 − CD19, BANK1 − LY9, POU2F2 − S1PR4 (sARI: 0.89, 0.58, 0.37, 0.32, 0.27) which are all B-cell immune signature genes and the bottom row contains the same for CD6 − CD8A, ARHGAP9 − BIN2, RASSF5 − BIN2, IL23A − CD96, CD6 − CD2 (sARI: 0.95, 0.91, 0.74, 0.71, 0.69) corresponding to T-cell immune signature genes. Corresponding plots of other spatially varying edges based on GEP for both B-cell and T-cell signature genes are provided in Figure S17 of the Supplementary Materials. Among the spatially varying edge plots in Figure 5, we observe a higher level of connectivity among the T-cell signature genes than the B cell signature genes. We identify ARHGAP9 − BIN2 (Figure 5H) has more local connectivity outside the tumor regions whereas IL23A − CD96 (Figure 5J) shows a constant pattern across the tissue and high local spatial connectivity in the tumor region than other regions.

**Figure 5.**
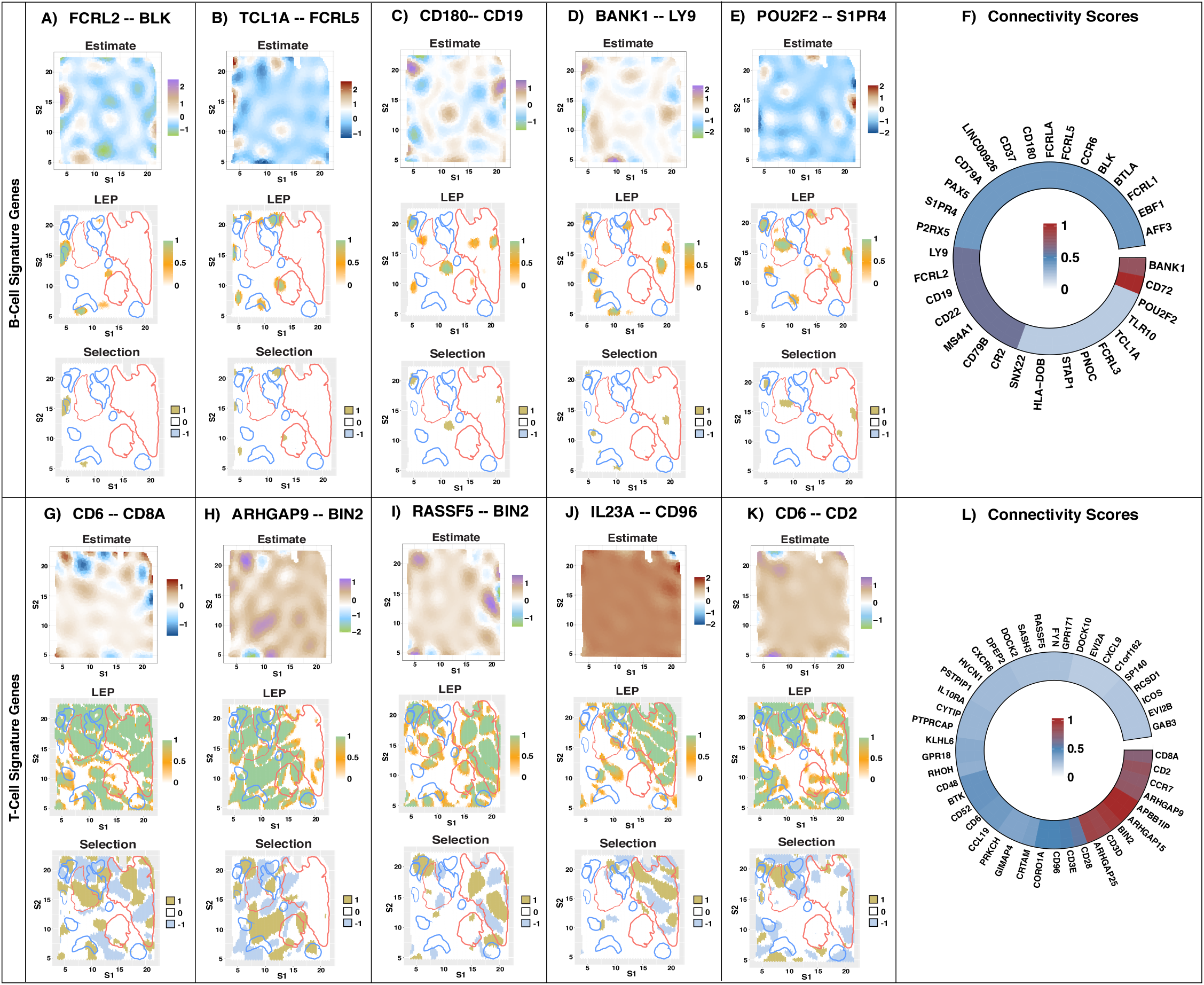
Spatially varying edges and connectivity scores for breast cancer: The Figure shows spatially varying edges for immune signature genes for B-cell (top row) and T-cell (bottom row). We provide spatially varying local edge probabilities and selection along with positive/negative interaction for each edge. The red and blue demarcation on the LEP and selection plot indicates tumor and normal region respectively. Rest of the region is considered as an intermediate region. The top row contains the spatially varying edges between FCRL2 − BLK, TCL1A − FCRL5, CD180 − CD19, BANK1 − LY9, POU2F2 − S1PR4 among the B-cell immune signature genes. The bottom row presents the spatially varying edges between CD6 − CD8A, ARHGAP9 − BIN2, RASSF5 − BIN2, IL23A − CD96, CD6 − CD2 among the T-cell immune signature genes. Figure 5F and 5L shows the circular heatmap of a matrix where each entry represents the number of connections of a immune signature gene w.r.t. B-cell and T-cell respectively. Color represents the gene connectivity levels (red, high; blue, medium; white low).

Figures 5F and 5L depict circular heatmaps representing a matrix in which each cell signifies the connectivity degree of an immune signature genes of B-cells and T-cells, respectively. In Figures 5F and 5L, the color gradient (red, high; blue, medium; white low) indicates the level of connectivity for genes. We filter out some of genes with low scaled connectivity degree (*<* 0.2) for the T-cell immune signature gene panel and scaled the connectivity degree in both Figure 5F and 5L within [0, 1] for comparison purposes. A circular heatmap containing all the genes of T-cell gene signature panel is provided in Figure S18 of the Supplementary Materials.

Based on our analysis of B-cell gene signature module, we identify CD19 has high connectivity degree (= 0.6). CD19 is a protein-coding gene that plays a critical role in maintaining the balance between immune activation among B-cells, and immune response regulation [Wang et al., 2012]. We observe that CD19 is connected with CD180 (Figure 5C) which is part of toll-like receptor family [Chiron et al., 2008] that recognizes pathogen-associated molecular patterns. CD180 activation leads to the production of proinflammatory cytokines, which are essential for innate immune responses [Edwards et al., 2023]. This gene-gene association is crucial for B-cell development, maturation, differentiation and part of critical signaling pathways for the tumor microenvironment of HER2+ breast cancer patients [Garaud et al., 2019]. Among T-cell immune gene signature panel, we observe spatially varying edges related to BIN2 and CD3D which are identified high connectivity degrees (0.93 and 0.87). BIN2 (Bridging Integrator 2) is a member of the BIN-Amphiphysin-Rvs (BAR) domain protein family and the interactions of BAR domain proteins, including BIN2, can interfere with tumor cell growth [Prendergast et al., 2009]. CD3D is identified as a prognostic biomarker, and high CD3D expression is linked to lower survival rates in breast cancer patients Zhu et al. [2021]. CD3D has a strong correlation with T-cells, suggesting that CD3D upregulation could enhance T-cell immune infiltration in the tumor microenvironment and boost antitumor immunity by activating T lymphocytes [Yang et al., 2020].

### 5.2 Prostate cancer

The ST data is acquired by performing biopsy of the tissue sectioned at a thickness of 5*µ*m from stage III adenocarcinoma of human prostate. The hematoxylin and eosin staining image of the human prostate cancer tissue is provided in 4D. The areas exhibiting dark staining are indicative of a potential tumor region, while the remaining portion can be categorized as intermediate and normal regions. Analogous to the breast cancer dataset, the tumor and normal spatial regions are distinguished by red and blue colored boundaries while the rest of the tissue section can be described as intermediate region. The prostate cancer data consists of *n* = 4, 371 spots and the spatial region annotations are obtained from the 10X Genomics website.

The number of B-cell and T-cell immune signature genes are 28 and 77 respectively. The networks of B-cell and T-cell signature genes are provided in Figure 4E and 4F. We use the same threshold from the breast cancer data analysis and select the genes with high scaled connectivity degree (*>* 0.5) for the network of T-cell immune signature genes. A network plot with all the gene for the prostate cancer tissue is provided in Figure S19 of the Supplementary Materials. We follow the analogous steps from the previous data analysis and take a closer look at the some of the spatially varying edges with high index value of ARI. Figure 6 shows the estimate, LEP and selection with positive/negative interaction for B-cell (top row of Figure 6: CD19 − CD37, FCRLA − CD37, POU2F2 − HLA-DOB, CD79B − AFF3, S1PR4 − FCRL5 with sARI values 0.74, 0.64, 0.43, 0.34, 0.31 respectively) and T-cell (bottom row of Figure 6: RGS18 − ARHGAP25, HVCN1 − ARHGAP9, HVCN1 − ARHGAP15, FYN − CXCR6, DOCK2 − CD28 with sARI values 0.96, 0.88, 0.67, 0.56, 0.53) signature genes. In Figure 6J, the spatial edge between FYN − CXCR6 indicated more activity in normal region than the tumor region but HVCN1 − ARHGAP9 (Figure 6H) shows a reverse pattern with high activity in the tumor region. Figure S20 in the Supplementary Materials displays the spatially varying edges with high GEP values of global connectivity along with their corresponding LEP and selection plots. We show the connectivity degree of both set of gene modules though a circular heatmap (Figure 6F and 6L) where each entry represents the number of connection of that particular gene. Using the same threshold, we exclude genes with low scaled connectivity degree (*<* 0.2) for the T-cell immune signature gene panel and provide the connectivity plot for all the T-cell signature genes in Figure S21 of the Supplementary Materials.

**Figure 6:**
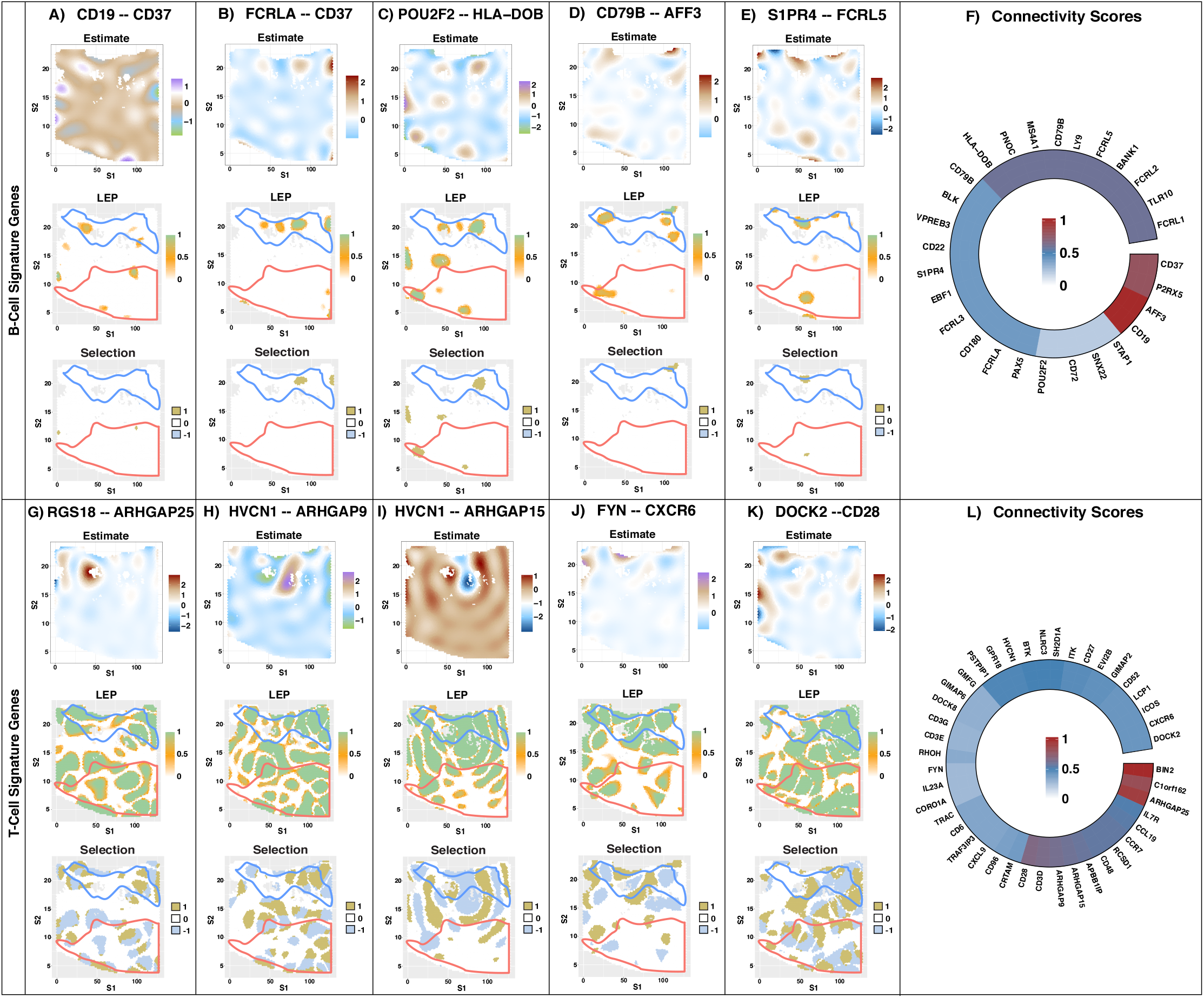
Spatially varying edges and connectivity scores for prostate cancer: The Figure displays spatially varying edges pertaining to immune signature genes associated with B-cells (top row) and T-cells (bottom row). We provide spatially varying local edge probabilities and selection information for each edge, including indications of positive or negative interactions. The red and blue colored markings on the LEP and selection plot signifies the tumor and normal regions, respectively, while the remaining region is regarded as an intermediate area. The top row contains the spatially varying edges between CD19 − CD37, FCRLA − CD37, POU2F2 − HLA-DOB, CD79B − AFF3, S1PR4 − FCRL5 amount the B-cell immune signature genes. The bottom row presents the spatially varying edges between RGS18 − ARHGAP25, HVCN1 − ARHGAP9, HVCN1 − ARHGAP15, FYN − CXCR6, DOCK2 − CD28 among the T-cell immune signature genes. Figure 6F and 6L shows the connectivity scores of B-cell and T-cell signature genes respectively. The color scheme indicates the intensity of gene connections, with red, blue, and white indicating high, medium, and low levels of connectivity respectively.

We detect AFF3 as hub gene (scaled connectivity degree = 1) for the B-cell immune gene signature panel. This gene is involved in RNA processing and plays a potential role in lymphocyte development and associated with tumorigenesis in prostate cancer [Fan et al., 2022]. Another hub gene POU2F2 plays a crucial role in both the proliferation and differentiation of B-cells into plasma cells by acting as a transcription regulator. Specifically, it activates immunoglobulin gene expression in mature B-cells, contributing significantly to the development of the immune system [Yang et al., 2021]. Multiple ARHGAP family genes are identified as hub genes (ARHGAP25, ARHGAP15, ARHGAP9) with scaled connectivity more than 0.6. These genes contribute to the modulation of variety of cellular functions such as cell structure, growth and motility. Genetic alteration of ARHGAP family genes initiates the cancer biogenesis [Chen et al., 2019]. In Figure 6, we observe more spatially varying significant patterns among the T-cell immune signature genes than B-cell ones which indicates T-cell activation to initiate anti-tumor response.

## 6 Conclusion and discussion

We propose a flexible `sGR` framework incorporating spatial information to estimate spatially-varying graphical models. `sGR` employs a spatial non-linear functional map between the spatial domain and spatially varying precision matrices enabling characterization of spatially varying edges through spatial conditional precision functions and identification of (local) spatial sub-regions where the function (edges) are significant using multiplicity controls. We demonstrate the efficacy of our method for estimation accuracy and graph structural recovery through rigorous simulation studies against existing methods. `sGR` is motivated by and applied to two ST data sets from breast and prostate cancer tissues to investigate gene regulatory networks in the tumor microenvironment. We estimate the spatially varying edges and infer genes with high values of degree connectivity. We identify several important hub genes from the CD (cluster of differentiation) and ARHGAP family among the T-cell immune signature genes in breast and prostate cancer tissue respectively. These hub genes could potentially help to develop T-CAR (T-cell chimeric antigen receptor) immunotherapy [Han et al., 2019] since T-cells can infiltrate tumor regions, and their interactions with cancer cells in the TME can influence cancer progression.

`sGR` employs a neighborhood selection approach that allows both for computational efficiency and sparse prior elicitation. For example, for B-cell signature genes for both datasets (*n* ≈ 4500, *p* ≈ 30 leading to 2 × 10^6^ possible edges) the `sGR` model takes around 2 and 8 hours for breast and prostate cancer data respectively to run on a high-computing cluster. We have employed a full MCMC algorithm for estimation, however alternate algorithms such as variational Bayes approaches could be explored to increase scalability.

As modern spatial technologies mature there are several applications and generalizations of `sGR` that are possible. The field of spatial-omics, which involves the high-throughput characterization of the transcriptome and proteome within the tissue context, is rapidly evolving. `sGR` can be used to study the spatially varying regulation pattern of any genomic features from spatial proteomics [Wu et al., 2022], and spatial metabolomics [Niehaus et al., 2019]. One could extend `sGR` to multiple samples (tissues) to model both intra- and inter-sample networks. Finally, integration of multiple types of spatial omics data, such as genomics, proteomics, and epigenomics require development of integrative spatial networks, as has been done for bulk sequencing data [Ha et al., 2021]. We leave these tasks for future explorations.

## SUPPLEMENTARY MATERIAL

**Title:** The Supplementary pdf file includes additional details on posterior computation, MCMC sampling, additional results of simulation studies and ST data analysis.

**R-package:** The R package is available at https://github.com/*****.

**Data set:** The breast and prostate cancer datasets are obtained from the 10X genomics.

