## Supplemental files for "Spatially varying graph estimation for spatial transcriptomics cancer data"

### Supplementary for "Spatial Graphical Regression Models for Spatial Transcriptomics Cancer Data"

May 4, 2025

In the Supplementary Materials, we provide detailed steps of posterior computation and algorithm related discussion in Section S1. Section S2 and Section S3 contains additional results from simulation study and spatial transcriptomics data analysis.

### S1 Methodological details

#### S1.1 Spatial kernel decomposition

Based on Mercer's theorem (Williams and Rasmussen, 2006), we perform Karhunen Loève expansion on SCPF i.e. $\beta_{ij}(s)$ and rewrite as

$$\begin{aligned}&\beta_{ij}(s) = \sum_{l=1}^{\infty} \varphi_l^{\beta}(s) u_{lij}^{\beta}, \text{ where } u_{lij}^{\beta} \sim N(0, \tau_{ij}^2 \psi_l^{\beta}) \text{ independently with } \psi_l^{\beta} > 0 \text{ s.t.} \\ &\boldsymbol{\mathcal{K}}_S^{\beta}(s, s') = \sum_{l=1}^{\infty} \psi_l^{\beta} \varphi_l^{\beta}(s) \varphi_l^{\beta}(s') \text{ and } \int \varphi_l^{\beta}(s) \varphi_{l'}^{\beta}(s) ds = 0 \ \forall \ l \neq l'.\end{aligned} \tag{S1}$$

This is an element wise expansion of the equation (4) of the manuscript. We truncate the infinite sum into $L_\beta$ terms such that $\sum_{l=1}^{L} \zeta_l^{\beta} / \sum_{l=1}^{\infty} \zeta_l^{\beta}$ is close to 1.

Next, we'll use Proposition 1 in Shi and Kang (2015) for the construction of $\psi_l$ and $\varphi_l(s)$ for equation (S1) in case of a spatial modified squared exponential function as discussed in the Section 3 of the manuscript. The closed form of eigen decomposition of MSE kernel can be obtained where eigen functions are expressed as function of Hermite polynomial. More specifically the eigen values and eigen functions can be written as

$$\begin{aligned}\psi_l^{\beta} &= \left(\frac{\pi}{A}\right)^n B^m, \\ \varphi_l^{\beta}(s) &= (2c)^{\frac{n}{4}} \exp(-c \mid\mid s \mid\mid_2^2) \prod_{q=1}^{2} H_{m_i}(\sqrt{2c} s_q),\end{aligned}$$

where ${}^{m+n-1}C_n < l \leq {}^{m+n}C_n$, $s = (s_1, s_2)$, $c = \sqrt{a^2 + 2ab}$, $A = a + b + c$ and $B = b/A$. The function $H_{m_i}(x)$ denotes normalized Hermit polynomial of non-negative order $\{m_i\}_{i=1}^n$ such that $\sum_{i=1}^n m_i = m$ and $H_k(x) = (2^k k! \sqrt{\pi})^{-1/2}(-1)^k \exp(x^2) \frac{d^k}{dx^k} \exp(-x^2)$. We use the same KL expansion (S1) on the spatial random effect $\eta_i(s)$ as well.

#### S1.2 Posterior computation

We briefly restate the spatially varying Gaussian graphical model and the spatial graphical regression (sGR) method before diving into the details of posterior computation. For notational purposes, we denote the spatial domain $\boldsymbol{\mathcal{S}}$ with $n$ spatial locations as $\boldsymbol{\mathcal{S}} = (s_1, \ldots, s_n)$ and the matrix from p-dimensional features is $\mathbf{Y}(\boldsymbol{\mathcal{S}}) = [\underline{Y}_1(\boldsymbol{\mathcal{S}}), \ldots, \underline{Y}_p(\boldsymbol{\mathcal{S}})] \in \mathbb{R}^{n \times p}$. The spatially varying Gaussian graphical model is provided as

$$\underline{Y}(s) \sim \text{MVN}\{\underline{\mu}(s), \boldsymbol{\Omega}(s)^{-1}\}, \quad \text{where} \;\; \boldsymbol{\Omega}(s) = (\omega_{ij}(s))_{p \times p}$$

where $\boldsymbol{\Omega}(s)$ is the spatially varying precision matrix for a given location $s \in \boldsymbol{\mathcal{S}}$. More details are provided in Section 2.2 of the manuscript. The undirected sGR is provided as

$$\begin{aligned}
&\underline{Y}_i(\boldsymbol{\mathcal{S}}) = \sum_{j \neq i}^{p} \underline{Y}_j(\boldsymbol{\mathcal{S}}) \underline{\beta}_{ij}(\boldsymbol{\mathcal{S}}) \gamma_{ij} + \underline{\eta}_i(\boldsymbol{\mathcal{S}}) + \underline{\epsilon}_i, \quad \underline{\epsilon}_i \sim N(0, \sigma^2_{\epsilon_i} \boldsymbol{I}_n), \qquad 1 \leq i \neq j \leq p, \\
&\underline{\beta}_{ij}(.) \sim \mathcal{GP}[0, \tau^2_{ij} \boldsymbol{\mathcal{K}}^{\beta}_S], \quad \gamma_{ij} \sim \text{Ber}(p_{ij}), \\
&\underline{\eta}_i(.) \sim \mathcal{GP}[0, \zeta^2_i \boldsymbol{\mathcal{K}}^{\eta}_S], \qquad \underline{\epsilon}_i \sim N(0, \sigma^2_{\epsilon_i} I_n).
\end{aligned}$$

Further details about the sGR model are discussed in Section 2.3 of the manuscript. We implement the method through a set parallel spatial regressions and derive the detailed steps of posterior computation for a fix node $i$. The likelihood can be expressed as

$$\begin{aligned}
L &= (2\pi\sigma^2_{\epsilon_i})^{-\frac{n}{2}} \exp\left[ -\frac{1}{2\sigma^2_{\epsilon_i}} \left\| \underline{Y}_i(\boldsymbol{\mathcal{S}}) - \sum_{j \neq i}^{p} \underline{Y}_j(\boldsymbol{\mathcal{S}}) \underline{\beta}_{ij}(\boldsymbol{\mathcal{S}}) \gamma_{ij} - \underline{\eta}_i(\boldsymbol{\mathcal{S}}) \right\|^2 \right] \\
&= (2\pi\sigma^2_{\epsilon_i})^{-\frac{n}{2}} \exp\left[ -\frac{1}{2\sigma^2_{\epsilon_i}} \left\| \underline{Y}_i(\boldsymbol{\mathcal{S}}) - \sum_{j \neq i}^{p} \gamma_{ij} \sum_{l=1}^{L} u^{\beta}_{lij} \underline{Y}_j(\boldsymbol{\mathcal{S}}) \underline{\varphi}^{\beta}_l(\boldsymbol{\mathcal{S}}) - \sum_{l=1}^{L} u^{\eta}_{li} \underline{\varphi}^{\eta}_l(\boldsymbol{\mathcal{S}}) \right\|^2 \right]
\end{aligned}$$

The expression for priors are provided as follows

$$\begin{aligned}
P = &\left[ \prod_{j \neq i}^{p} \prod_{l=1}^{L} N(u^{\beta}_{lij} \mid 0, \zeta^{\beta}_l) \right] \times \left[ \prod_{l=1}^{L} N(u^{\eta}_{li} \mid 0, \zeta^{\eta}_l) \right] \times \\
&\text{Inv-Gamma}(\sigma^2_{\epsilon_i} \mid a_{\epsilon_i}, b_{\epsilon_i}) \times \left[ \prod_{j \neq i}^{p} \prod_{l=1}^{L} \text{Ber}(\gamma_{ij} \mid p_{ij} = 0.5) \right].
\end{aligned}$$

A simple algebraic derivation which will help us to get conditional update equations are provided below

$$
\begin{aligned}
&\left|\left|\underline{Y}_i(\boldsymbol{\mathcal{S}}) - \sum_{j\neq i}^{p} \underline{Y}_j(\boldsymbol{\mathcal{S}})\underline{\beta}_{ij}(\boldsymbol{\mathcal{S}})\gamma_{ij} - \underline{\eta}_i(\boldsymbol{\mathcal{S}})\right|\right|^2 \\
&= \left|\left|\underline{Y}_i(\boldsymbol{\mathcal{S}}) - \sum_{k\neq\{i,j\}}^{p} \underline{Y}_k(\boldsymbol{\mathcal{S}})\underline{\beta}_{ik}(\boldsymbol{\mathcal{S}})\gamma_{ik} - \underline{Y}_j(\boldsymbol{\mathcal{S}})\underline{\beta}_{ij}(\boldsymbol{\mathcal{S}})\gamma_{ij} - \underline{\eta}_i(\boldsymbol{\mathcal{S}})\right|\right|^2 \\
&= \left|\left|\underline{O}_{-j}(\boldsymbol{\mathcal{S}}) - \gamma_{ij}\underline{Y}_j(\boldsymbol{\mathcal{S}})\underline{\beta}_{ij}(\boldsymbol{\mathcal{S}})\right|\right|^2 = \left|\left|\underline{O}_{-j}(\boldsymbol{\mathcal{S}}) - \gamma_{ij}\underline{Y}_j(\boldsymbol{\mathcal{S}})\sum_{l=1}^{L} u^{\beta}_{lij}\underline{\varphi}^{\beta}_{l}(\boldsymbol{\mathcal{S}})\right|\right|^2 \\
&= \left|\left|\underline{O}_{-j}(\boldsymbol{\mathcal{S}}) - \sum_{l'\neq l}^{L} u^{\beta}_{l'ij}\gamma_{ij}\underline{Y}_j(\boldsymbol{\mathcal{S}})\underline{\varphi}^{\beta}_{l'}(\boldsymbol{\mathcal{S}}) - u_{lij}\gamma_{ij}\underline{Y}_j(\boldsymbol{\mathcal{S}})\underline{\varphi}^{\beta}_{l}(\boldsymbol{\mathcal{S}})\right|\right|^2 = \left|\left|\underline{O}^{-l}_{-j}(\boldsymbol{\mathcal{S}}) - u^{\beta}_{lij}\gamma_{ij}\underline{Y}_j(\boldsymbol{\mathcal{S}})\underline{\varphi}^{\beta}_{l}(\boldsymbol{\mathcal{S}})\right|\right|^2 \\
&= \left|\left|\underline{O}^{-l}_{-j}(\boldsymbol{\mathcal{S}}) - u^{\beta}_{lij}\underline{\tilde{Y}}_{ij,l}(\boldsymbol{\mathcal{S}})\right|\right|^2 = || \underline{\tilde{Y}}_{ij,l}(\boldsymbol{\mathcal{S}}) ||^2 \, (u^{\beta}_{lij})^2 - 2u^{\beta}_{lij}(\underline{\tilde{Y}}^{T}_{ij,l}(\boldsymbol{\mathcal{S}})\underline{O}^{-l}_{-j}(\boldsymbol{\mathcal{S}})) - || \underline{O}^{-l}_{-j}(\boldsymbol{\mathcal{S}}) ||^2,
\end{aligned}
$$

$$
\underline{O}_{-j}(\boldsymbol{\mathcal{S}}) = \underline{Y}_i(\boldsymbol{\mathcal{S}}) - \sum_{k\neq\{i,j\}}^{p} \underline{Y}_k(\boldsymbol{\mathcal{S}})\underline{\beta}_{ik}(\boldsymbol{\mathcal{S}})\gamma_{ik} - \underline{\eta}_i(\boldsymbol{\mathcal{S}}), \quad \underline{O}^{-l}_{-j}(\boldsymbol{\mathcal{S}}) = \underline{O}_{-j}(\boldsymbol{\mathcal{S}}) - \sum_{l'\neq l}^{L} u^{\beta}_{l'ij}\gamma_{ij}\underline{Y}_j(\boldsymbol{\mathcal{S}})\underline{\varphi}^{\beta}_{l'}(\boldsymbol{\mathcal{S}}),
$$

$$
\underline{\tilde{Y}}_{ij,l}(\boldsymbol{\mathcal{S}}) = \gamma_{ij}\underline{Y}_j(\boldsymbol{\mathcal{S}})\underline{\varphi}^{\beta}_{l}(\boldsymbol{\mathcal{S}}).
$$

Combining the likelihood, prior and previously mentioned derivations, we derive the conditional update of the parameters as discussed below.

- **Conditional update of** $u^{\beta}_{lij}$:

$$
\begin{aligned}
&q(u^{\beta}_{lij} \mid -) \propto \exp\left[-\frac{1}{2\sigma^2_{\epsilon_i}}\{|| \underline{\tilde{Y}}_{ij,l}(\boldsymbol{\mathcal{S}}) ||^2 \, (u^{\beta}_{lij})^2 - 2u^{\beta}_{lij}(\underline{\tilde{Y}}^{\mathrm{T}}_{ij,l}(\boldsymbol{\mathcal{S}})\underline{O}^{-l}_{-j}(\boldsymbol{\mathcal{S}}))\}\right] \\
&\qquad\qquad \times \exp\left[-\frac{1}{2\zeta^{\beta}_{l}}(u^{\beta}_{lij})^2\right], \\
&(u^{\beta}_{lij} \mid -) \sim N(\mu^{\beta}_{u_{lij}}, \sigma^{\beta}_{u_{lij}}), \\
&\mu^{\beta}_{u_{lij}} = (\sigma^{\beta}_{u_{lij}})^{-1}[\underline{\tilde{Y}}^{\mathrm{T}}_{ij,l}(\boldsymbol{\mathcal{S}})\underline{O}^{-l}_{-j}(\boldsymbol{\mathcal{S}})], \quad \sigma^{\beta}_{u_{lij}} = \left[\frac{|| \underline{\tilde{Y}}_{ij,l}(\boldsymbol{\mathcal{S}}) ||^2}{\sigma^2_{\epsilon_i}} + \frac{1}{\zeta^{\beta}_{l}}\right]^{-1}.
\end{aligned}
$$

- **Conditional update of $u^{\eta}_{li}$:**

$$\left|\left|\underset{\sim}{Y}_i(\boldsymbol{\mathcal{S}}) - \sum_{j\neq i}^{p} \underset{\sim}{Y}_j(\boldsymbol{\mathcal{S}})\underset{\sim}{\beta}_{ij}(\boldsymbol{\mathcal{S}})\gamma_{ij} - \underset{\sim}{\eta}_i(\boldsymbol{\mathcal{S}})\right|\right|^2 = \left|\left|\underset{\sim}{Y}_i(\boldsymbol{\mathcal{S}}) - \sum_{j\neq i}^{p} \underset{\sim}{Y}_j(\boldsymbol{\mathcal{S}})\underset{\sim}{\beta}_{ij}(\boldsymbol{\mathcal{S}})\gamma_{ij} - \sum_{l=1}^{L} u^{\eta}_{li}\underset{\sim}{\varphi}^{\eta}_l(\boldsymbol{\mathcal{S}})\right|\right|^2$$

$$= \left|\left|\underset{\sim}{\tilde{O}}^{-l}_i(\boldsymbol{\mathcal{S}}) - u^{\eta}_{li}\underset{\sim}{\varphi}^{\eta}_l(\boldsymbol{\mathcal{S}})\right|\right|^2 \text{ where } \underset{\sim}{\tilde{O}}^{-l}_i(\boldsymbol{\mathcal{S}}) = \underset{\sim}{Y}_i(\boldsymbol{\mathcal{S}}) - \sum_{j\neq i}^{p} \underset{\sim}{Y}_j(\boldsymbol{\mathcal{S}})\underset{\sim}{\beta}_{ij}(\boldsymbol{\mathcal{S}})\gamma_{ij} - \sum_{l'\neq l}^{L} u^{\eta}_{l'i}\underset{\sim}{\varphi}^{\eta}_{l'}(\boldsymbol{\mathcal{S}})$$

$$= || \underset{\sim}{\varphi}^{\eta}_l(\boldsymbol{\mathcal{S}}) ||^2 (u^{\eta}_{li})^2 - 2u^{\eta}_{li}[\{\underset{\sim}{\varphi}^{\eta}_l(\boldsymbol{\mathcal{S}})\}^T \underset{\sim}{\tilde{O}}^{-l}_i(\boldsymbol{\mathcal{S}})] - || \underset{\sim}{\tilde{O}}^{-l}_i(\boldsymbol{\mathcal{S}}) ||^2 .$$

Based on the previous derivations, we have the conditional update of $u^{\eta}_{li}$ as follows

$$q(u^{\eta}_{li} \mid -) \propto \exp\left[-\frac{1}{2\sigma^2_{\epsilon_i}}\{|| \underset{\sim}{\varphi}^{\eta}_l(\boldsymbol{\mathcal{S}}) ||^2 (u^{\eta}_{li})^2 - 2u^{\eta}_{li}[\{\underset{\sim}{\varphi}^{\eta}_l(\boldsymbol{\mathcal{S}})\}^T \underset{\sim}{\tilde{O}}^{-l}_i(\boldsymbol{\mathcal{S}})]\}\right] \times \exp\left[-\frac{1}{2\zeta^{\eta}_l}(u^{\eta}_{li})^2\right],$$

$$(u^{\eta}_{li} \mid -) \sim N(\mu^{\eta}_{u_{li}}, \sigma^{\eta}_{u_{li}}),$$

$$\mu^{\eta}_{u_{li}} = (\sigma^{\eta}_{u_{li}})^{-1}[\{\underset{\sim}{\varphi}^{\eta}_l(\boldsymbol{\mathcal{S}})\}^T \underset{\sim}{\tilde{O}}^{-l}_i(\boldsymbol{\mathcal{S}})], \quad \sigma^{\eta}_{u_{li}} = \left[\frac{|| \underset{\sim}{\varphi}^{\eta}_l(\boldsymbol{\mathcal{S}}) ||^2}{\sigma^2_{\epsilon_i}} + \frac{1}{\zeta^{\eta}_l}\right]^{-1}.$$

- **Conditional update of $\sigma^2_{\epsilon_i}$:**

$$(\sigma^2_i \mid -) \sim \text{Inv-Gamma}\left[\frac{n}{2} + a_{\epsilon_i}, b_{\epsilon_i} + \frac{1}{2}\left|\left|\underset{\sim}{Y}_i(\boldsymbol{\mathcal{S}}) - \sum_{j\neq i}^{p} \underset{\sim}{Y}_j(\boldsymbol{\mathcal{S}})\underset{\sim}{\beta}_{ij}(\boldsymbol{\mathcal{S}})\gamma_{ij} - \underset{\sim}{\eta}_i(\boldsymbol{\mathcal{S}})\right|\right|^2\right].$$

- **Conditional update of $\gamma_{ij}$:**

$$(\gamma_{ij} \mid -) \sim \text{Ber}(\tilde{p}_{ij}) \text{ s.t. } \tilde{p}_{ij} = \frac{c_{ij}}{c_{ij} + d_{ij}},$$

$$c_{ij} = p_{ij} L(\underset{\sim}{\tilde{\beta}}^{*}_{ij}(\boldsymbol{\mathcal{S}})),$$

$$d_{ij} = (1 - p_{ij}) L(\underset{\sim}{\tilde{\beta}}^{**}_{ij}(\boldsymbol{\mathcal{S}})),$$

$$\underset{\sim}{\tilde{\beta}}^{*}_{ik}(\boldsymbol{\mathcal{S}}) = \begin{cases} \underset{\sim}{\beta}_{ij}(\boldsymbol{\mathcal{S}}) & k = j \\ \underset{\sim}{\tilde{\beta}}_{ik}(\boldsymbol{\mathcal{S}}) & k \neq j \end{cases}, \quad \underset{\sim}{\tilde{\beta}}^{**}_{ik}(\boldsymbol{\mathcal{S}}) = \begin{cases} 0 & k = j \\ \underset{\sim}{\tilde{\beta}}_{ik}(\boldsymbol{\mathcal{S}}) & k \neq j. \end{cases}$$

$L(\underset{\sim}{\tilde{\beta}}^{*}_{ij}(\boldsymbol{\mathcal{S}}))$ and $L(\underset{\sim}{\tilde{\beta}}^{**}_{ij}(\boldsymbol{\mathcal{S}}))$ denotes the likelihood containing $\underset{\sim}{\tilde{\beta}}^{*}_{ij}(\boldsymbol{\mathcal{S}})$ and $\underset{\sim}{\tilde{\beta}}^{**}_{ij}(\boldsymbol{\mathcal{S}})$ where $\underset{\sim}{\tilde{\beta}}_{ij}(\boldsymbol{\mathcal{S}}) = \underset{\sim}{\beta}_{ij}(\boldsymbol{\mathcal{S}})\gamma_{ij}$. This completes our computational details of our MCMC algorithm.

#### S1.3 Positive definite property

For a spatial location, we assume that the true precision matrix is diagonal dominant. The positive definite property of the final estimator is ensured through diagonal dominance which

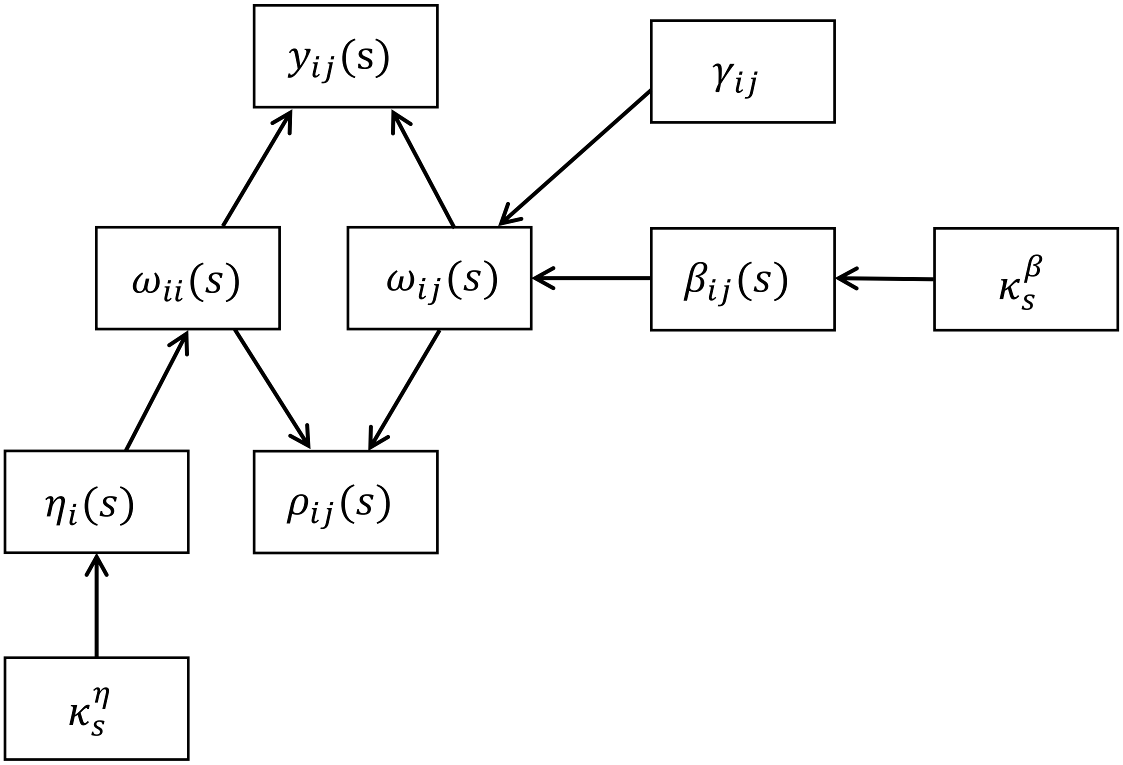


Figure S1: A schematic representation of the `sGR` model.

holds true if

$$\Rightarrow \mid \omega_{ii} \mid \geq \sum_{j \neq i} \mid \omega_{ij}(s) \mid = \sum_{j \neq i} \mid -\omega_{ii}\beta_{ij}(s) \mid$$

$$\Rightarrow \mid \omega_{ii} \mid \geq \sum_{j \neq i} \mid -\omega_{ii}(\sum_{l=1}^{L_\beta} u_{lij}^\beta \varphi_l^\beta(s)) \mid$$

$$\Rightarrow 1 \geq \sum_{j \neq i} \mid \sum_{l=1}^{L_\beta} u_{lij}^\beta \varphi_l^\beta(s) \mid$$

We also make an assumption that $\mathbf{\Phi}^\beta(\boldsymbol{\mathcal{S}})$ is within known ranges i.e. $\mid\mid \mathbf{\Phi}^\beta(\boldsymbol{\mathcal{S}}) \mid\mid_\infty = a < \infty$. We can also write the following steps of inequality.

$$\sum_{j \neq i} \mid \sum_{l=1}^{L_\beta} u_{lij}^\beta \varphi_l^\beta(s) \mid \leq \sum_{j \neq i} \sum_{l=1}^{L_\beta} \mid u_{lij}^\beta \varphi_l^\beta(s) \mid \leq a \sum_{j \neq i} \sum_{l=1}^{L_\beta} \mid u_{lij}^\beta \mid = a \mid\mid U_{i\cdot}^\beta \mid\mid_1$$

Here $U^{\beta}_{i\bullet}$ is a $(p-1)\times L_{\beta}$ dimensional matrix which contains basis coefficients. Combining both inequalities, we have

$$\frac{\sum_{j\neq i} \mid \sum_{l=1}^{L_\beta} u^{\beta}_{lij}\varphi^{\beta}_{l}(s) \mid}{a \mid\mid U^{\beta}_{i\bullet} \mid\mid_1} \leq 1$$

Therefore, we can obtain the diagonal dominance matrix as a final estimate if we rescale the basis coefficients as $\underline{U}^{\beta}_{ij} = \underline{U}^{\beta}_{ij}/(a \mid\mid U^{\beta}_{i\bullet} \mid\mid_1)$.

#### S1.4 Glocal-local probability scores

In equation (5) of the manuscript, we define the $100(1-\alpha)\%$ joint credible bands for SCPF $\beta_{\bullet}(s)$ as

$$\Pr\{L_{\bullet}(s) \leq \beta_{\bullet}(s) \leq U_{\bullet}(s), \forall s \in \boldsymbol{\mathcal{S}} \subset \mathbb{R}^2\} \geq (1-\alpha), \tag{S2}$$

where $L_{\bullet}(s)$ and $U_{\bullet}(s)$ the lower and upper bound respectively. We adapt the steps from Ruppert et al. (2003) under a spatial setting to further express an interval satisfying equation (S2) can be rewritten as

$$I_{\bullet,\alpha}(s) = \hat{\beta}_{\bullet}(s) \pm q_{(1-\alpha)}[\widehat{\text{St.Dev}}\{\hat{\beta}_{\bullet}(s)\}],$$

where $\hat{\beta}_{\bullet}(s)$ and $\widehat{\text{St.Dev}}\{\hat{\beta}_{\bullet}(s)\}$ are the mean and standard deviation for a given $s$ taken over all the MCMC samples. The variable $q_{(1-\alpha)}$ denotes the $(1-\alpha)$ quantile over $M$ posterior samples taken over the quantity

$$Z^{(m)}_{\bullet} = \max_{S\in\boldsymbol{\mathcal{S}}} \left| \frac{\beta^{(m)}_{\bullet}(s) - \hat{\beta}_{\bullet}(s)}{\widehat{\text{St.Dev}}\{\hat{\beta}_{\bullet}(s)\}} \right|.$$

These joint credible bands do not require a pre-specified $\delta$-fold intensity change like BFDR and provides strong experiment-wise control for multiple testing. Further steps to construct local probability scores is discussed in Section 3.2 of the manuscript.

### S2 Additional results from simulation study

In this section, we provide additional simulation study related details. The graph structural recovery rates are defined as TPR = TP/(TP + FN), FPR = FP/(FP + TN), FDR = FP / (FP + TP) and

$$\text{MCC} = \frac{\text{TP} \times \text{TN} - \text{FP} \times \text{FN}}{\sqrt{(\text{TP+FP})(\text{TP+FN})(\text{TN+FP})(\text{TN+FN})}}$$

where TP, FP, TN, and FN denotes true positives, false positives, true negatives, and false negatives. For the undirected settings discussed in the Section 4 of the manuscript, the `sGR` method yields high values of TPR (`sGR`: 0.98, GraphR: 0.90, Go CART: 0.87, Penalized GR: 0.76, Glasso: 0.72) but at the cost of low values of FDR (`sGR`: 0.065, GraphR: 0.183, Go CART: 0.204, Penalized GR: 0.456, Glasso: 0.471) and FPR (`sGR`: 0.004, GraphR: 0.005, Go CART: 0.008, Penalized GR: 0.011, Glasso: 0.018).

The functional form of 12 SCPFs from Figure 3A is provided below

$$\beta_{42,24} = 2\tanh(s_1^2 + \tanh^{-1}(s_2)), \quad \beta_{63,21} = s_1^3 + s_2^3,$$
$$\beta_{49,20} = 2 \mid s_1^2 + s_2^3 \mid -2, \quad \beta_{19,13} = 3\tanh(s_1 - s_2^3),$$
$$\beta_{28,8} = s_2(s_1^2 + 0.5), \quad \beta_{72,2} = s_2(s_1 + 0.5),$$
$$\beta_{43,7} = (2(s_1 + s_2)) * exp(-0.5 * (s_1 - s_2)), \quad \beta_{43,22} = \tanh(s_1) - \tanh(s_2),$$
$$\beta_{8,6} = 2 \mid s_1 + s_2^3 \mid -2, \quad \beta_{67,2} = s_1 + s_2,$$
$$\beta_{54,11} = s_1 + s_2^2 + s_1^3 + s_2^4, \quad \beta_{67,17} = s_1 + exp(s_2) - 1.5.$$

In Figure S2, we provide the estimate, LEP and selection of rest of the 6 SCPFs out of 12 from Figure 3. Figure S3 − S14 shows true, estiamte, LEP and selection of corresponding SCPFs for signal-to-niose ratio = $\{1, 3, 5\}$. Table S1 lists all the functional forms of SCPFs.

We evaluate the frequentist properties of SGM in case of estimation accuracy and structural recovery with varying spatial locations and fixed number of nodes. To this end, we define the following norm $c_n^\beta = \frac{1}{n}\sum_{j<i}^{N_{edge}} \| \beta_{ij} - \hat{\beta}_{ij} \|_2$ where $N_{edge}$ denotes the total number of edges. Figure S15A shows the boxplot of $c_n^\beta$ across 50 replicates for different values of $n = \{1000, 2000, 3000, 4000, 5000\}$. We observe a decreasing pattern in the boxplot with increasing values of $n$ which indicates the consistency of our method. We use the ROC curves and the area under the ROC curve to compare the graph structual recovery performance for directed spatial graphical models across different settings of $(n, p)$ values. To summarize the performance across spatial locations and replicates, we calculate the true positive rates (TPR) and the false positive rates (FPR) by averaging the sensitivity and specificity of the directed spatial graphical model across replicates. These TPR and FPR values are used to develop ROC curve and AUC values. Figure S15B displays the increasing pattern in the ROC curves with higher values of $n$ which suggests the increasing behaviour of our method in terms of graph structural recovery with growing values of $n$. In summary, the directed `sGR` model setting performs better in terms of graph estimation and structural recovery with increasing number of spatial locations.

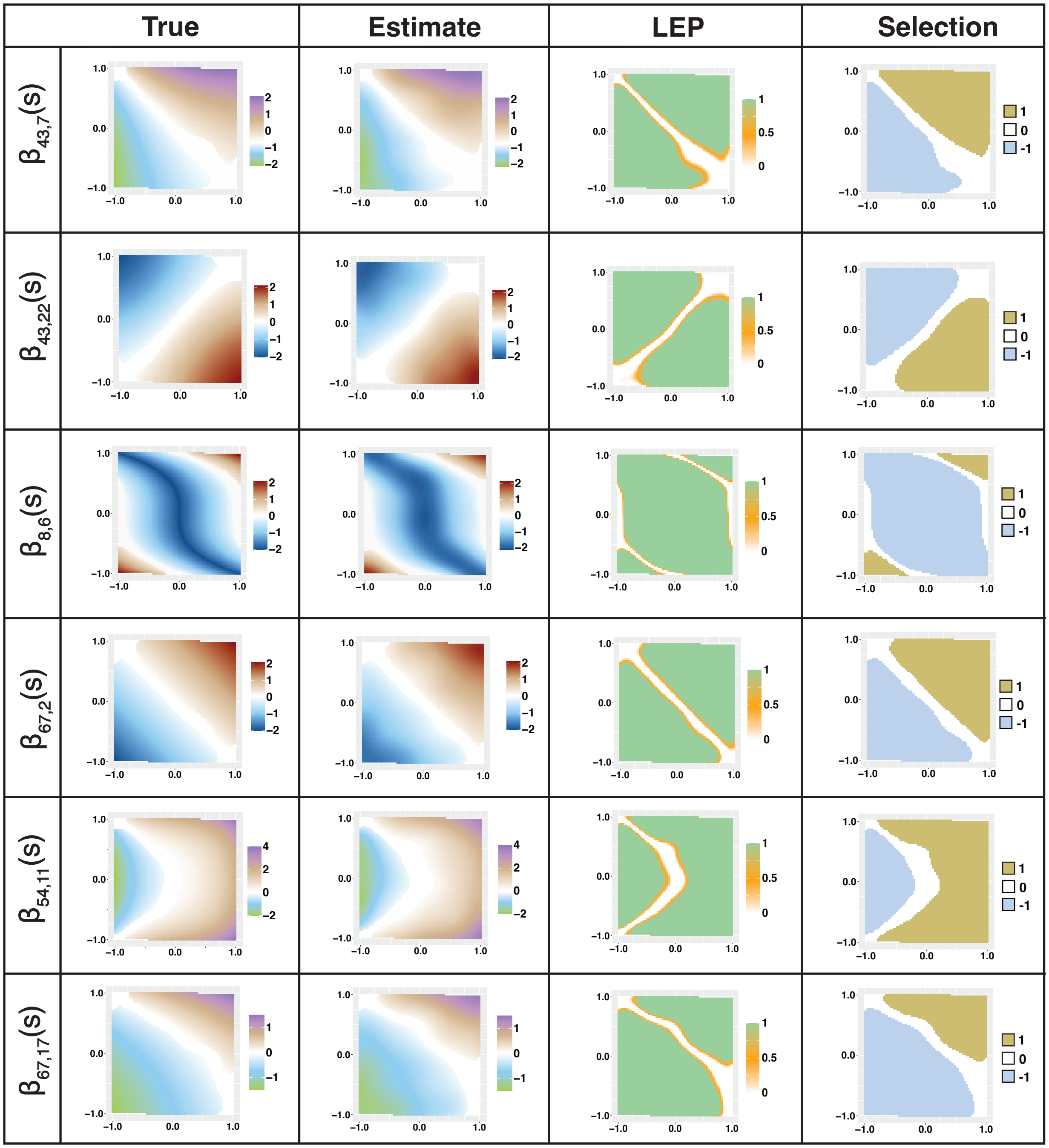


Figure S2: This Figure shows another set of 6 spatially varying edges ($\beta_{43,7}$, $\beta_{43,22}$, $\beta_{8,6}$, $\beta_{67,2}$, $\beta_{54,11}$, and $\beta_{67,17}$ out of 12 based on Figure 3. The column of the plot shows the true, estimate, LEP and selection of the SCPFs respectively.

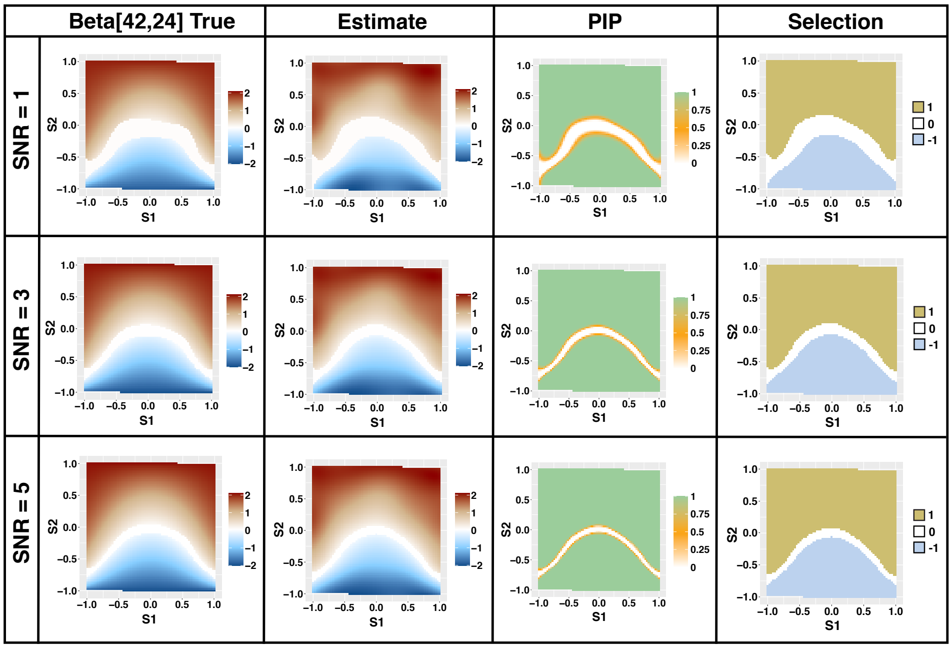

Figure S3: Estimation and selection of $\beta_{42,24}(s) = 2\tanh(s_1^2 + \tanh^{-1}(s_2))$ under directed spatial graphical model with $(n, p) = (5000, 100)$ and SNR$= \{1, 3, 5\}$.

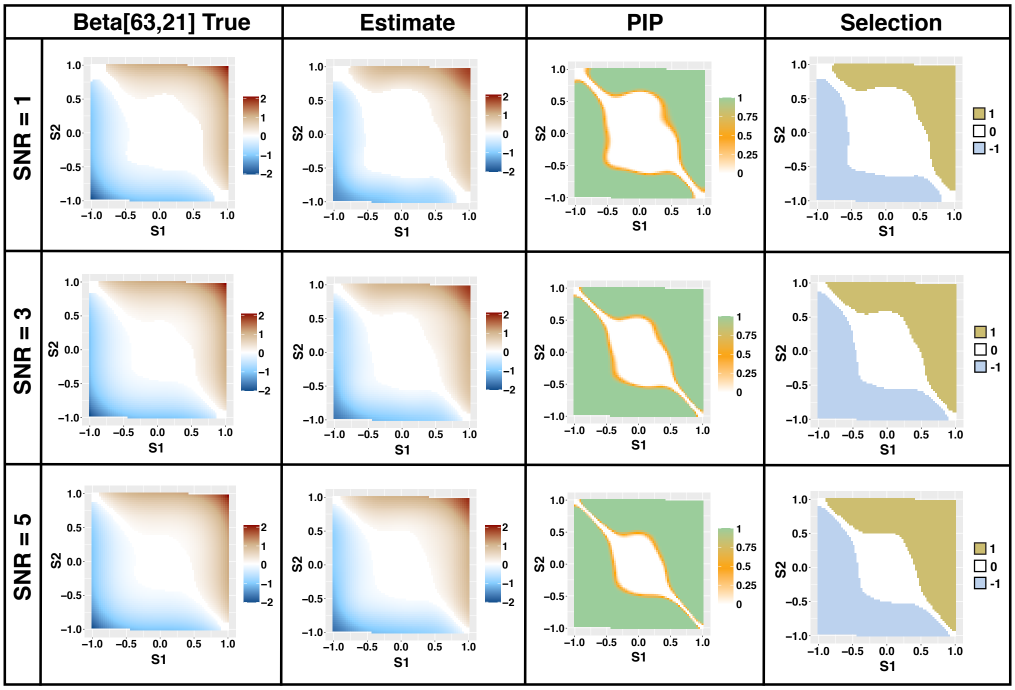

Figure S4: Estimation and selection of $\beta_{63,21}(s) = s_1^3 + s_2^3$ under directed spatial graphical model with $(n, p) = (5000, 100)$ and SNR$= \{1, 3, 5\}$.

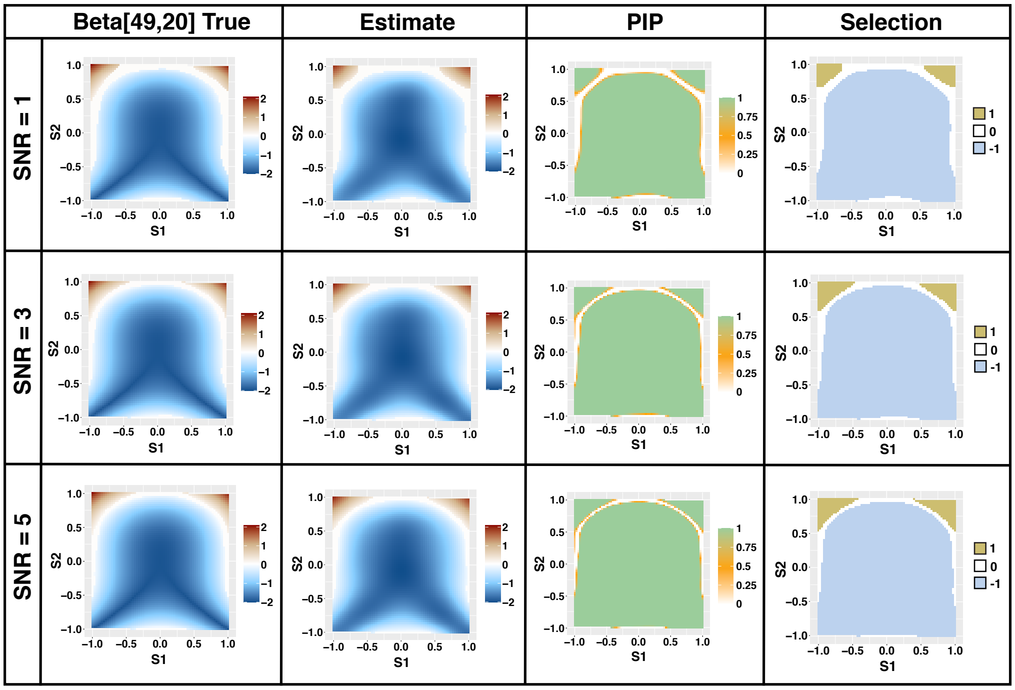


Figure S5: Estimation and selection of $\beta_{49,20}(s) = 2 \mid (s_1^2 + s_2^3) \mid -2$ under directed spatial graphical model with $(n, p) = (5000, 100)$ and SNR$= \{1, 3, 5\}$.

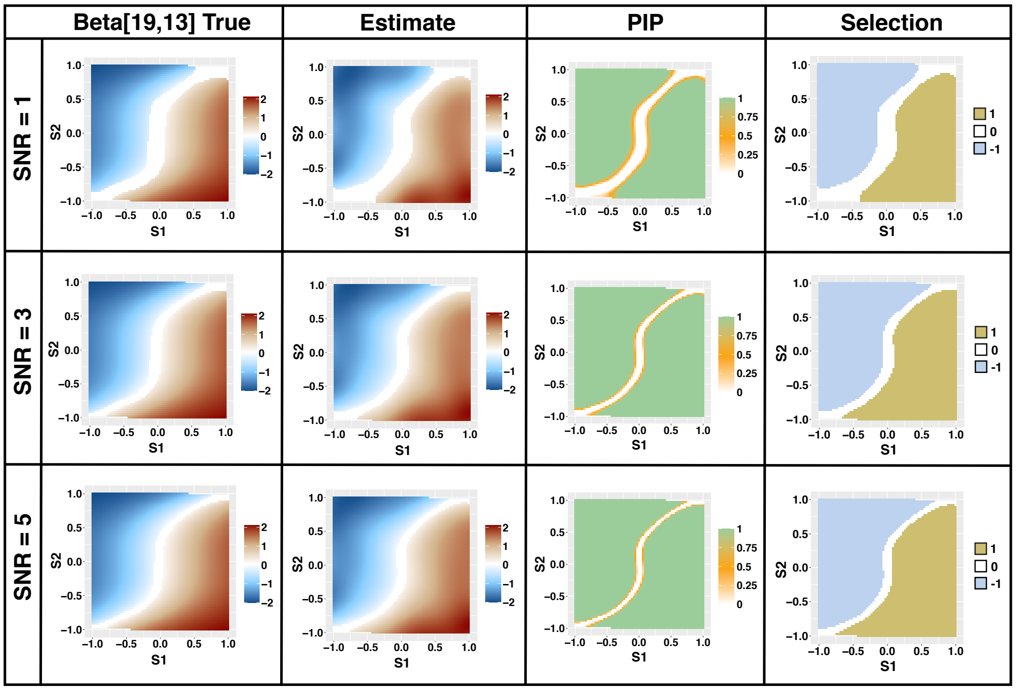


Figure S6: Estimation and selection of $\beta_{19,13}(s) = 3\tanh(s_1 - s_2^3)$ under directed spatial graphical model with $(n, p) = (5000, 100)$ and SNR$= \{1, 3, 5\}$.

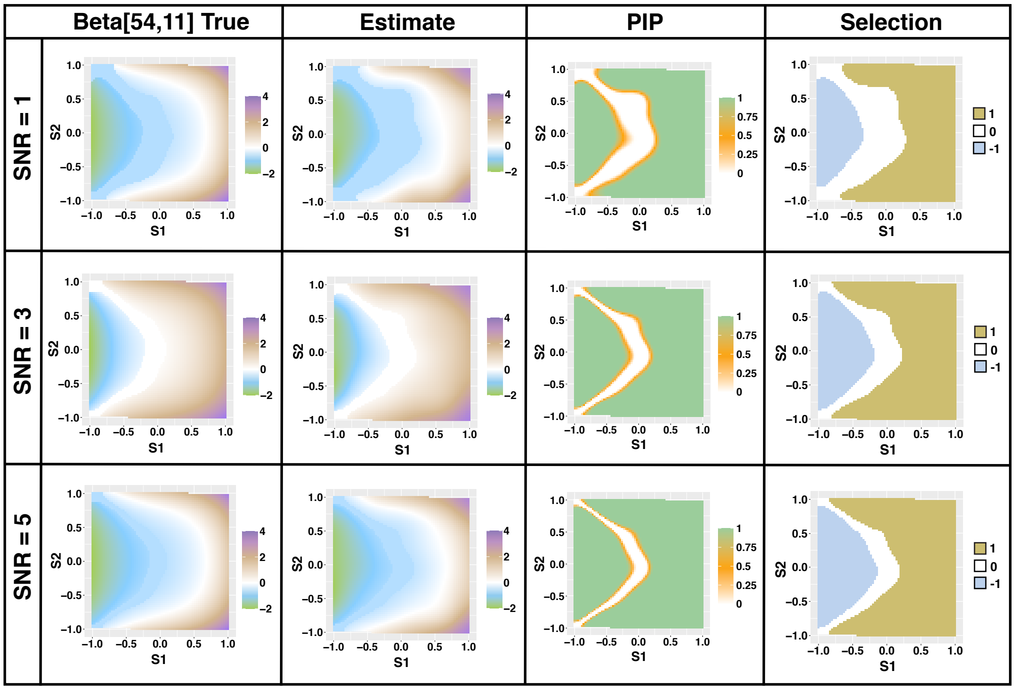


Figure S7: Estimation and selection of $\beta_{54,11}(s) = s_1 + s_2^2 + s_1^3 + s_2^4$ under directed spatial graphical model with $(n, p) = (5000, 100)$ and SNR$= \{1, 3, 5\}$.

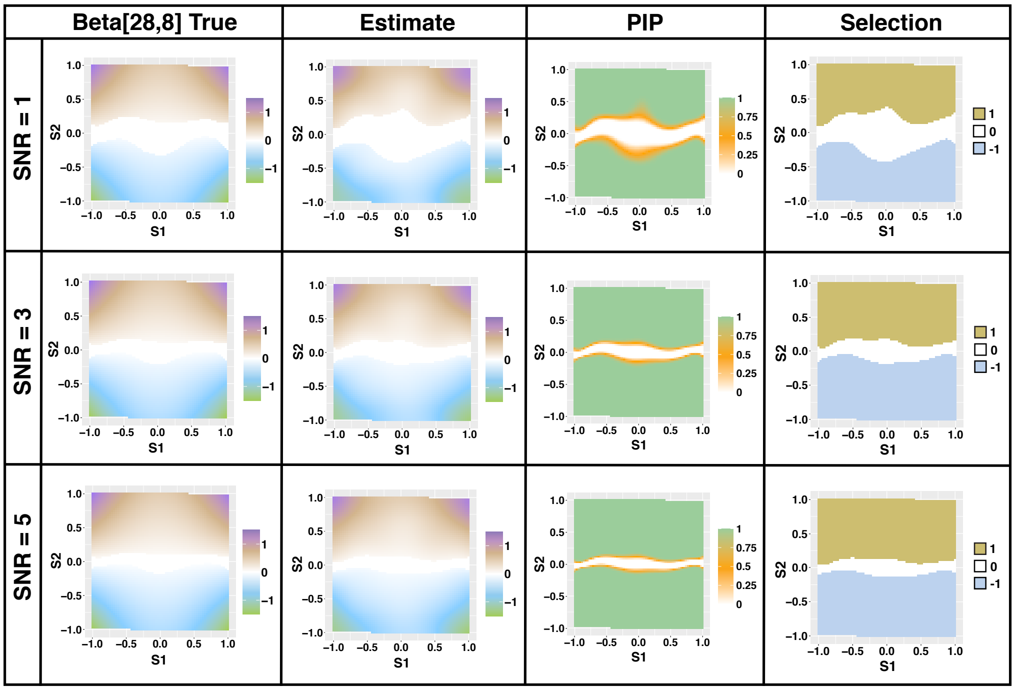


Figure S8: Estimation and selection of $\beta_{28,8}(s) = s_2 * (s_1^2 + 0.5)$ under directed spatial graphical model with $(n, p) = (5000, 100)$ and SNR$= \{1, 3, 5\}$.

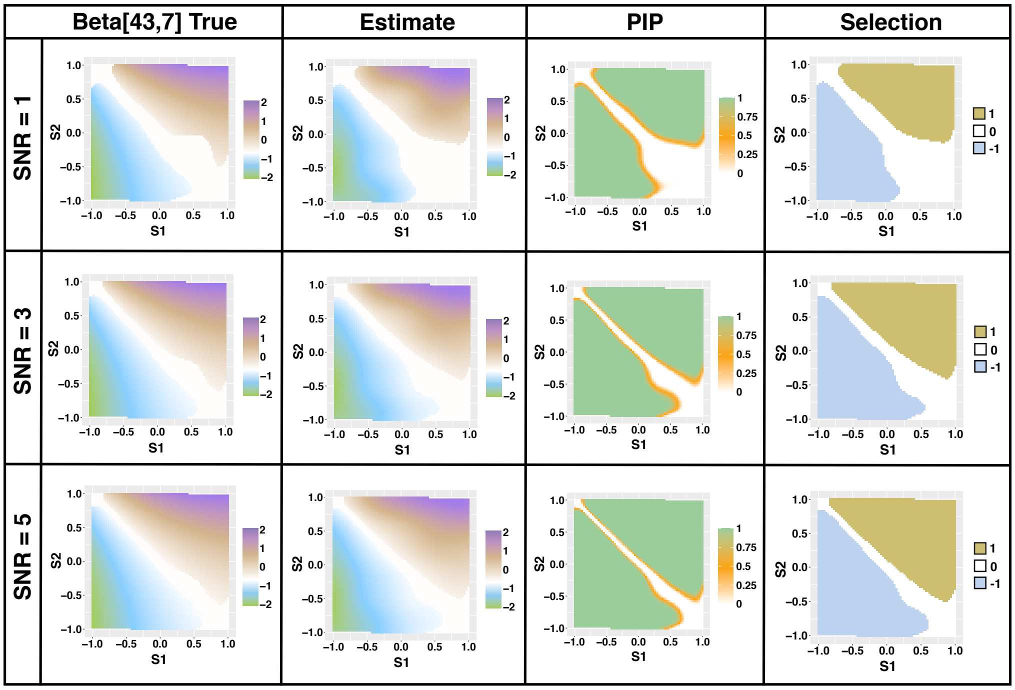


Figure S9: Estimation and selection of $\beta_{43,7}(s) = 2(s_1 + s_2) * \exp(-0.5 * (s_1 - s_2)$ under directed spatial graphical model with $(n, p) = (5000, 100)$ and SNR$= \{1, 3, 5\}$.

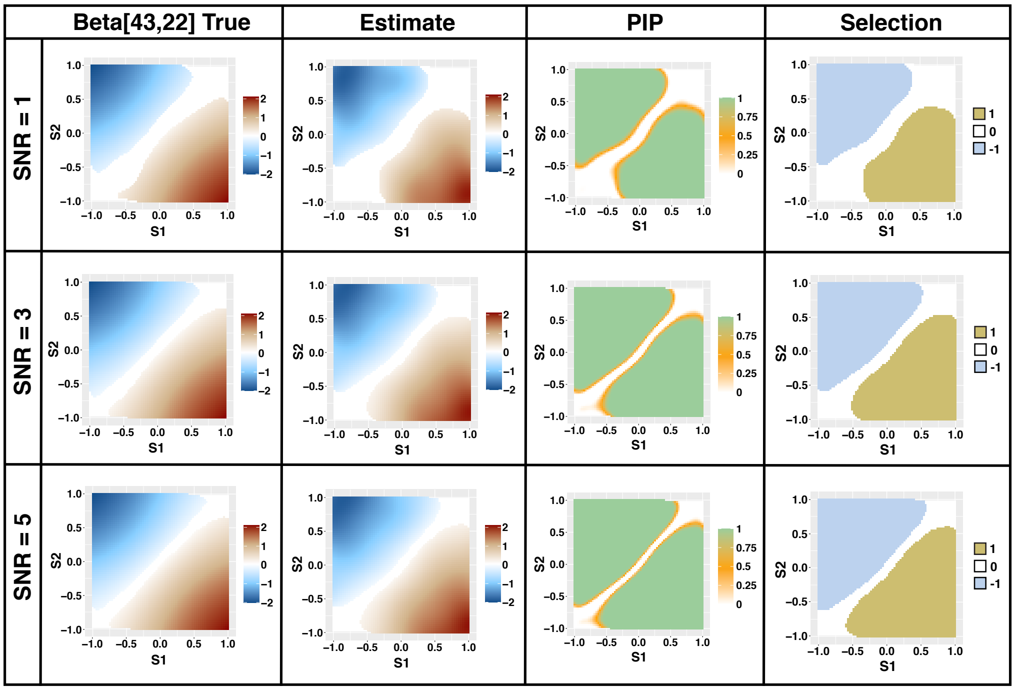


Figure S10: Estimation and selection of $\beta_{43,22}(s) = \tanh(s_1) - \tanh(s_2)$ under directed spatial graphical model with $(n, p) = (5000, 100)$ and SNR$= \{1, 3, 5\}$.

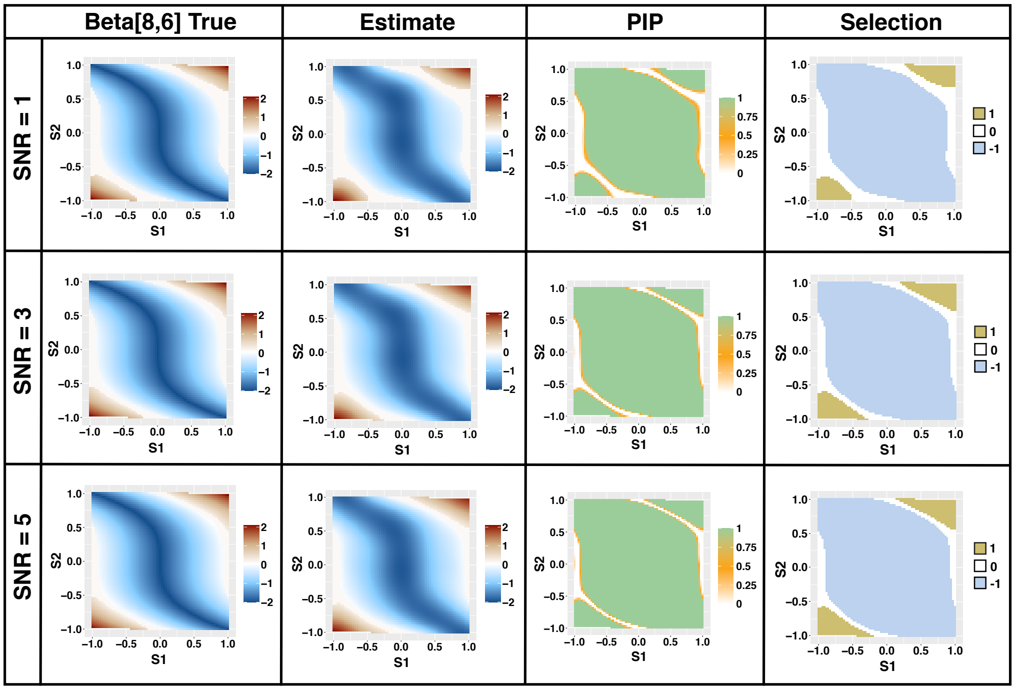


Figure S11: Estimation and selection of $\beta_{8,6}(s) = 2 \mid (s_1 + s_2^3) \mid -2$ under directed spatial graphical model with $(n, p) = (5000, 100)$ and SNR$= \{1, 3, 5\}$.

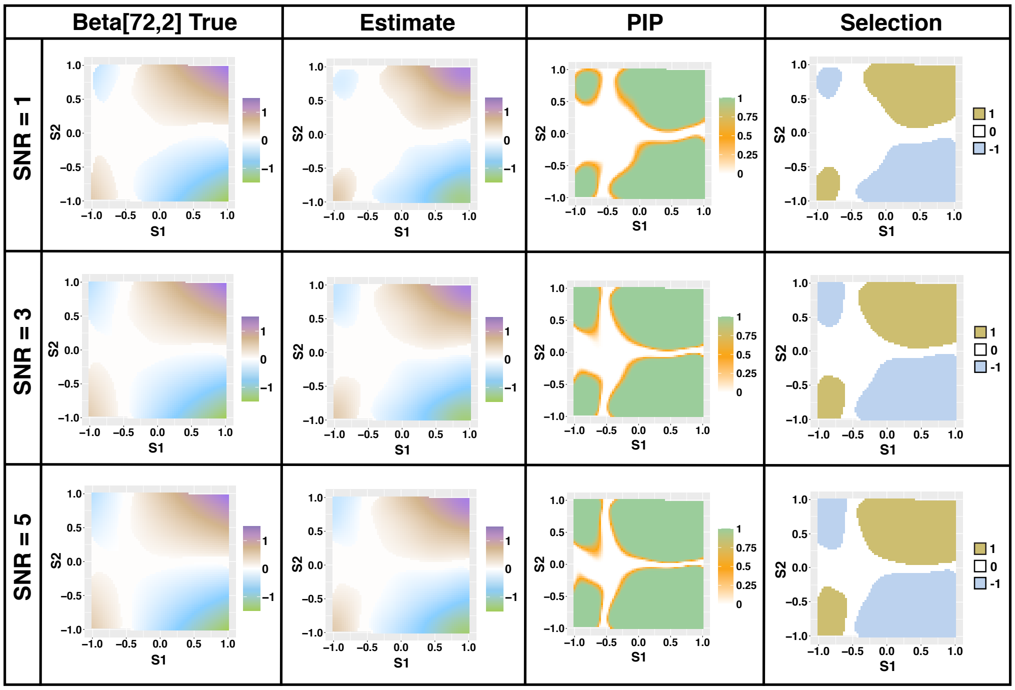


Figure S12: Estimation and selection of $\beta_{72,2}(s) = s_2 * (s_1 + 0.5)$ under directed spatial graphical model with $(n, p) = (5000, 100)$ and SNR$= \{1, 3, 5\}$.

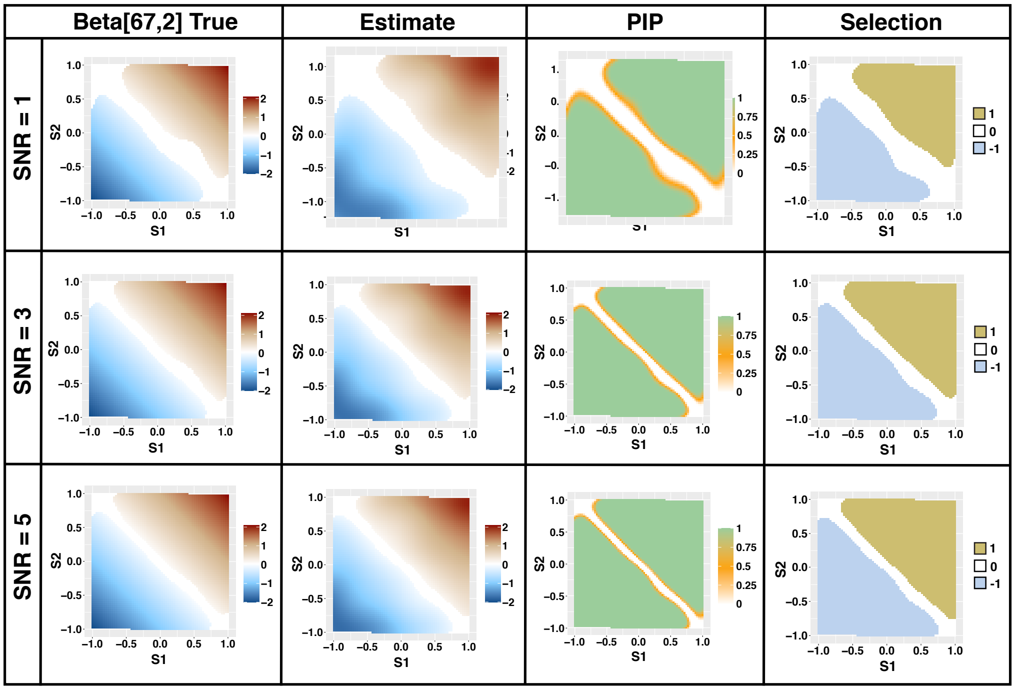


Figure S13: Estimation and selection of $\beta_{67,2}(s) = s_1 + s_2$ under directed spatial graphical model with $(n, p) = (5000, 100)$ and SNR$= \{1, 3, 5\}$.

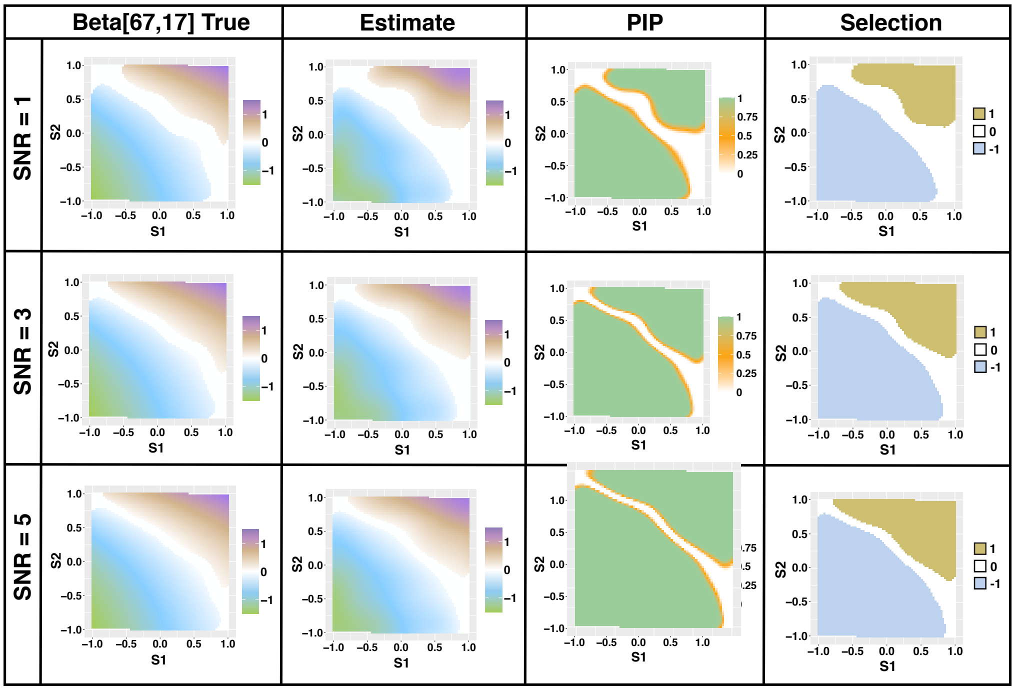


Figure S14: Estimation and selection of $\beta_{67,17}(s) = s_1 + \exp(s_2) - 1.5$ under directed spatial graphical model with $(n, p) = (5000, 100)$ and SNR$= \{1, 3, 5\}$.

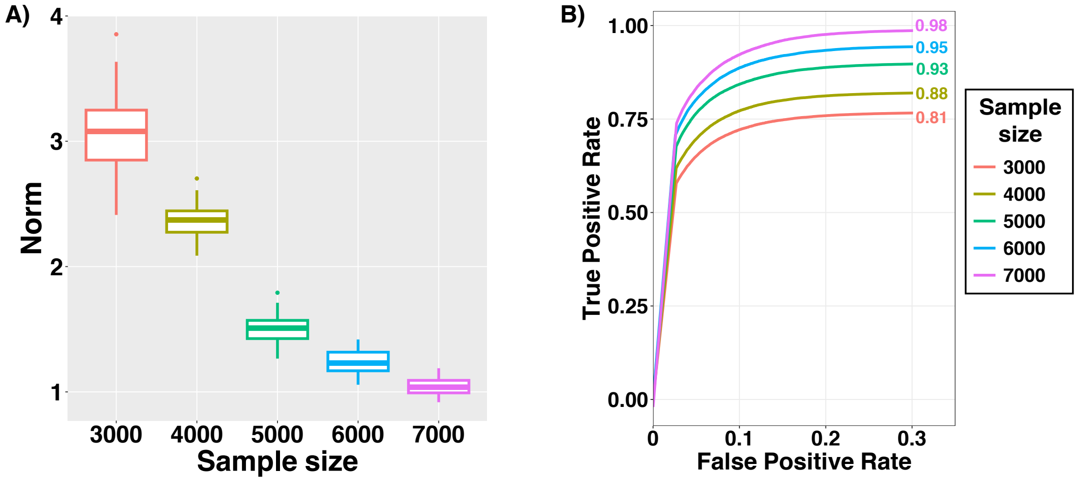


Figure S15: Performance of directed sGR model. Figure S15A shows the boxplot of $c_n^{\beta}$ across 50 replicates which summarizes the consistency in case of estimating spatially varying edges. Figure S15B indicates graph structural recovery pattern with ROC curves for increasing values of $n = \{3000, 4000, 5000, 6000, 7000\}$ and fix $p = 100$, SNR$= 3$.

| | | | |
|---|---|---|---|
| $S_1^3 + S_2^3$ | $\exp(S_1^2 + S_2^2)$ | $\log(S_1^4 + 1) + \exp(S_2)$ | $S_1 + S_2$ |
| $(S_1^2 + S_2^2 < 1.5)$ | $\exp(S_1) + \exp(S_2^2)$ | $S_1 + S_2^3$ | $\exp(S_1) + \exp(S_2) + 1$ |
| $\exp(\cos(S_1 - S_2))$ | $\log(S_1^2 + 1) - S_1^2 + 2$ | $\exp(-5(S_2 - 1.5\sin(\pi * \lvert S_1\rvert) + 1.0)^2) + 1$ | $\lvert S_1\rvert + \lvert S_2\rvert$ |
| $S_1 + S_2^2$ | $S_1^4 + S_2$ | $S_1 + \exp(S_2) - 1.5$ | $\exp(\cos(S_1 + S_2))$ |
| $\exp(\cos(S_1)) + \exp(\sin(S_2))$ | $\cos(S_1)^2 + S_2 + 1$ | $\exp(-0.5 * (S_1^2 + S_2^2))$ | $(2 * (S_1 + S_2)) * \exp(-0.5 * (S_1 - S_2))$ |
| $0.5 * (\exp(S_1) + \exp(S_2)) + 1$ | $0.5 * (\exp(S_1) - \exp(S_2))$ | $\log(S_1^2 + 1 + \mathrm{sqrt}(S_2^2 + 1)) + 0.25$ | $\log(S_1^2 + 5 - \mathrm{sqrt}(S_2^2 + 1)) + 0.5$ |
| $\exp(\sin(S_1^2)) + \cos(S_2)$ | $\exp(\sin(S_2^2)) - \cos(S_1)$ | $\lvert S_1\rvert - \lvert S_2\rvert$ | $S_1^3 + 4 - \mathrm{sqrt}(S_2 + 6)$ |
| $S_1^2 + 4 - sqrt(S_2 + 6)$ | $S_1 - S_2^2 + S_1^3$ | $S_1 + S_2^2 + S_1^3$ | $S_1 + S_2^2 - S_1^3 + 0.25$ |
| $S_1 - S_2^2 - S_1^3 - 0.5$ | $S_1 + S_2^2 + S_1^3 + S_2^4$ | $S_1 - S_2^2 + S_1^3 - S_2^4$ | $(-S_1) + S_2^2 - S_1^3 + S_2^4$ |
| $S_1 - (S_2 + 1)^2 - S_1^3 - S_2^4 + 2.5$ | $(S_1 + 0.5) * S_2$ | $(S_1^2 + 0.5) * (S_2)$ | $(S_1 + 0.5) * (S_2 - 1)$ |
| $(S_1^2 + 1.5) * (S_2^2 - 1) + 1$ | $(S_1^2 + 0.5) * (S_2^2 + 0.5)$ | $(S_1^2 + 0.25) * (S_2^3 - 1) + 1$ | $(S_1 + 0.5) * (S_2^4 + 0.5)$ |
| $(S_1 - 0.5) * (S_2^4 + 0.5)$ | $(S_1 + 1.5) * (S_2^4 - 1.5)$ | $(S_1 - 2) * (S_2^4 - 1.5)$ | $exp(-5 * (S_2^2 - 1.5 * sin(pi * abs(S_1)) + 1.0)^2) + 1$ |
| $\lvert S_1^3 + 1\rvert - \lvert S_2 + 1\rvert$ | $\lvert S_1^2\rvert + \lvert S_2\rvert + 0.5$ | $\lvert S_1^4 + 0.5\rvert - \lvert S_2^3 + 1\rvert$ | $2\lvert S_1 + S_2\rvert - 2$ |
| $2\lvert S_1^2 + S_2\rvert - 2$ | $2\lvert S_1 + S_2^3\rvert - 2$ | $2\lvert S_1^2 + S_2^3\rvert - 2$ | $2\lvert S_1^2 - S_2\rvert - 2$ |
| $2\lvert S_1^3 - S_2\rvert - 2$ | $\lvert (S_1 + S_2)^3\rvert - 2$ | $2^{(}S_1 + S_2)$ | $\exp(S_1 + \log_{10}(S_2^2 + 1))$ |
| $\exp(S_1^2 + \log_{10}(S_2^2 + 1))$ | $\exp(S_1 + \log_{10}(S_2^4 + 1))$ | $\exp(S_1^2 - \log_{10}(S_2^2 + 1))$ | $2 * tan^{-1}(S_1 + S_2)$ |
| $2 * tan^{-1}(S_1 - S_2)$ | $2 * \mathrm{cospi}(S_1^2 + S_2^2)$ | $-(S_1 + S_2) + 0.25 * (1.5 * S_1 + S_2)^2$ | $5 + (S_1 + S_2) - (S_1 + S_2)^2 + (S_1 + S_2)^3$ |
| $2 * tan^{-1}(S_1^2 + S_2^2)$ | $2 * tan^{-1}(S_1^3 + S_2^2)$ | $20 * \mathrm{dnorm}(S_1, 2, 10) + 20 * \mathrm{dnorm}(S_2, 2, 30) + 2$ | $20 * \mathrm{dnorm}(S_1, 2, 10) - 20 * \mathrm{dnorm}(S_2, 2, 30) + 3$ |
| $5 * \mathrm{dgamma}(S_1 + S_2, 3, 2)$ | $\sinh(S_1) - \sin(S_2)$ | $\sinh(S_1) - \sinh(S_2)$ | $\sinh(S_1) + \sinh(S_2)$ |
| $\sinh(S_1) + \sin(S_2)$ | $\cosh(S_1) + \cosh(S_2)$ | $\cosh(S_1 + S_2)$ | $3 * \tanh(S_1 + S_2)$ |
| $\tanh(S_1) + \tanh(S_2)$ | $\tanh(S_1) - \tanh(S_2)$ | $\cosh(S_1) + \sinh(S_2)$ | $\tanh(S_1^3) + \tanh(S_2^3)$ |
| $\tanh(S_1 + \tanh^{-1}(S_2))$ | $2 * \tanh(S_1^2 + \tanh^{-1}(S_2))$ | $\tanh(S_1 + S_2^3)$ | $2 * \tanh(\sinh(S_1) + \tanh^{-1}(S_2^3))$ |
| $3 * \tanh(S_1 - S_2^3)$ | $\tanh(S_1) - \tanh(S_2^3)$ | $2 * \tanh(S_1) - \tanh(S_2^2)$ | $2 * \sinh(S_1) + \cosh(S_2^2)$ |
| $(\lvert S_1\rvert + \lvert S_2\rvert < 1.5)$ | $(\lvert S_1\rvert + \lvert S_2\rvert < 1.25)$ | $(\lvert S_1\rvert + \lvert S_2\rvert^3 < 1.4)$ | $(\lvert S_1\rvert^4 + \lvert S_2\rvert^4 < 1)$ |
| $\exp(\cos(S_1)) + \exp(\sin(S_2)) < 3$ | $-(\lvert S_1\rvert + \lvert S_2\rvert < 1.25)$ | $-(\exp(\cos(S_1)) + \exp(\sin(S_2)) < 3)$ | $S_1 + S_2^4$ |

Table S1: List of SCPFs for the simulation settings.

### S3 Applications

#### S3.1 Preprocessing steps

Both data sets are publicly available from the 10X genomics website (breast cancer data and prostate cancer data). The breast cancer tissue sample contains 36601 genes and 4727 spatial locations and the prostate cancer tissue sample has 17943 genes and 4371 spatial locations. We focus on the B cell and T cell immune signature genes (Nirmal et al., 2018) for both dataset. We find (33, 79) genes in breast cancer tissue and (28,77) genes in prostate cancer tissue related (B, T) cell immune signature genes. We apply the PQLseq (Sun et al., 2019) algorithm to obtain the latent gene expressions which follow a Gaussian distribution. We implement the spatial graphical regression method on these latent gene expressions and scaled spatial coordinates while implementing the method.

#### S3.2 Additional results for breast cancer tissue

In this section, we provide the additional real data analysis results for the application of the spatial graphical regression method (Section 2.3) to the spatial transcriptomics data from the breast cancer tissue. The scaled ARI (sARI) is defined as sARI $=$ ARI/c where $c = ([100 * \text{ARI}_{\text{max}}] + 1)/100$. Figure S16 the network of all T cell immune signature genes based on their global posterior inclusion probability. In Figure S17, we display the spatial edges with high values of global spatial connectivity from B cell (top row: PAX5 $-$ CD79B, BLK $-$ BTLA, S1PR4 $-$ FCRL1, BTLA $-$ CCR6 and HLA-DOB $-$ CD19) and and T cell (bottom row: CD28 $-$ CD2, ITGAL $-$ IL7R, SH2D1A $-$ CD96, FLI1 $-$ CD28 and TBC1D10C $-$ GPR18). We present the spatially varying edge estimates along with spatially varying local-PIP, and positive/negative interaction plot. The PIP and selection plots are demarcated in red and blue, which symbolize the tumor and normal regions, respectively. Among the spatially varying edges, we observe high level of significant interaction among the T cell immune signature genes than the B cell genes. In top row of Figure S17, we can observe that the spatially varying edge S1PR4 $-$ FCRL1 is interacting in the normal regions whereas the same edge is inactive in the tumor region. Figure S18 represents the circular heatmap of connectivity matrix of all T cell immune signature genes where red, blue and white color denotes high, mid and low level of connectivity.

#### S3.3 Additional results for prostate cancer tissue

In this section, we present more comprehensive real data analysis results from applying the spatial graphical regression method (refer to Section 2.3) to the spatial transcriptomics data of prostate cancer tissue. The network of all T cell immune signature genes, based on their

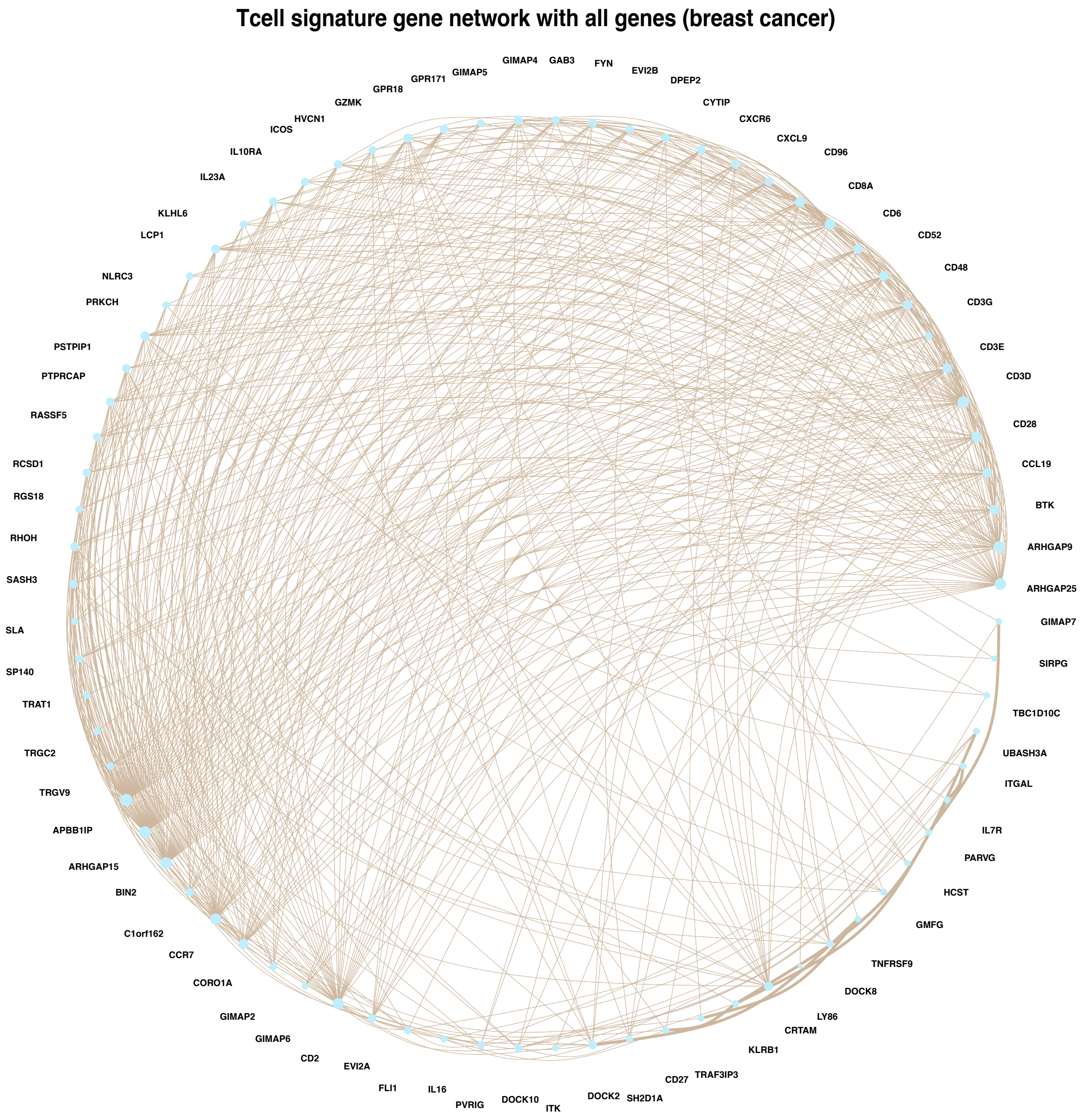


Figure S16: Global spatial connectivity network for T cell immune signature genes where all genes are included for breast cancer tissue.

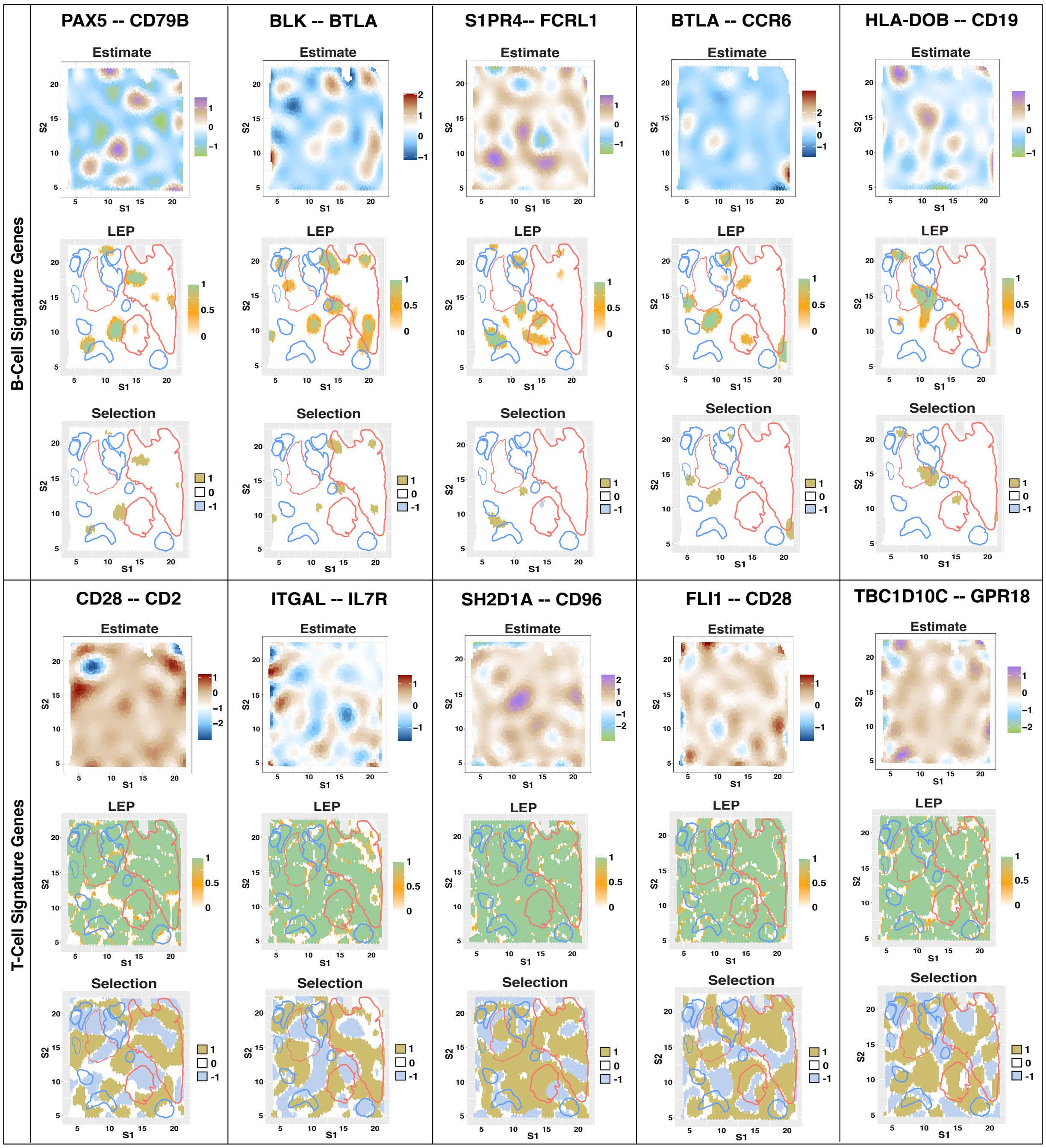


Figure S17: In case of the breast cancer tissue, the Figure shows spatially varying edges for immune signature genes for B cell (top row) and T cell (bottom row) based on their global posterior inclusion probability. We provide the estimate, spatially varying PIP and selection. The red and blue demarcation on the PIP and selection plot indicates tumor and normal region respectively. Rest of the region is considered as an intermediate region. The top row contains the spatially varying edges between PAX5 – CD79B, BLK – BTLA, S1PR4 – FCRL1, BTLA – CCR6 and HLA-DOB – CD19 among the B cell immune signature genes. The bottom row presents the spatially varying edges between CD28 – CD2, ITGAL – IL7R, SH2D1A – CD96, FLI1 – CD28 and TBC1D10C – GPR18 among the T cell immune signature genes.

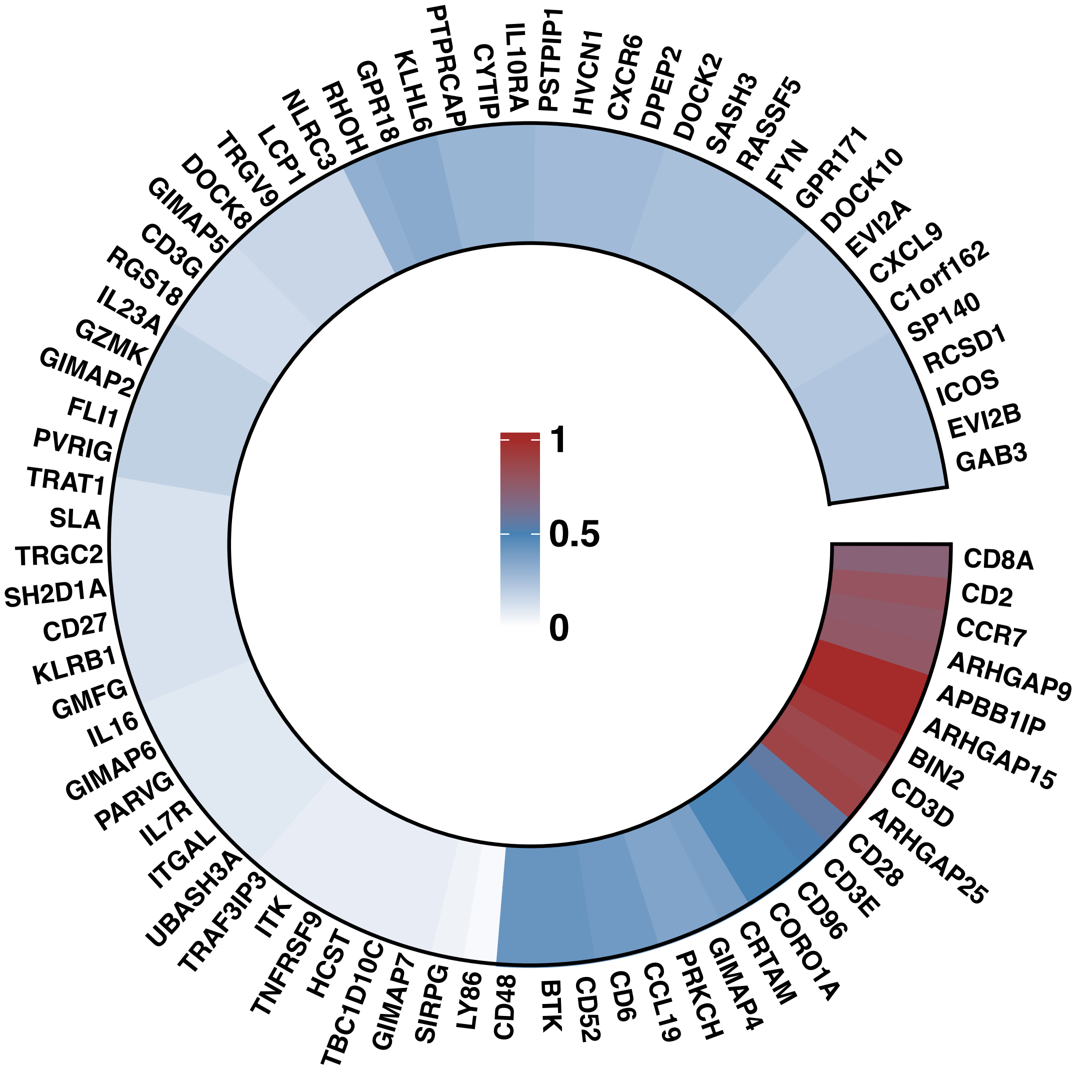


Figure S18: The Figure shows the circular heatmap of a matrix where each entry represents the number of connections of a immune signature gene w.r.t. T cell. Here we considered all the Tcell immune signature genes for the breast cancer tissue.

global posterior inclusion probabilities, is depicted in Figure S19. Figure S20 displays the spatial edges that have high global spatial connectivity values from both B cell (top row: VPREB3 − BANK1, CD180 − CD22, CD79A − EBF1, HLA-DOB − MS4A1, and FCRL5 − CD79A) and T cell (bottom row: CD3E − CXCR6, CYTIP − LY86, GMFG − CD52, PSTPIP1 − GPR171, and TRAC − CXCL9). Alongside these, we provide the spatially varying local-PIP, and the positive/negative interaction plot. We find that the spatially varying interaction between T cell immune signature genes are higher than the interaction between B cell immune signature genes. In the bottom row of Figure S20, the spatial edges CYTIP − LY86 and GMFG − CD52 are more active in the normal and transition region than the rumor region. Figure S21 depicts a circular heatmap of the connectivity matrix for all T cell immune signature genes. The colors red, blue, and white indicate high, medium, and low connectivity levels, respectively.

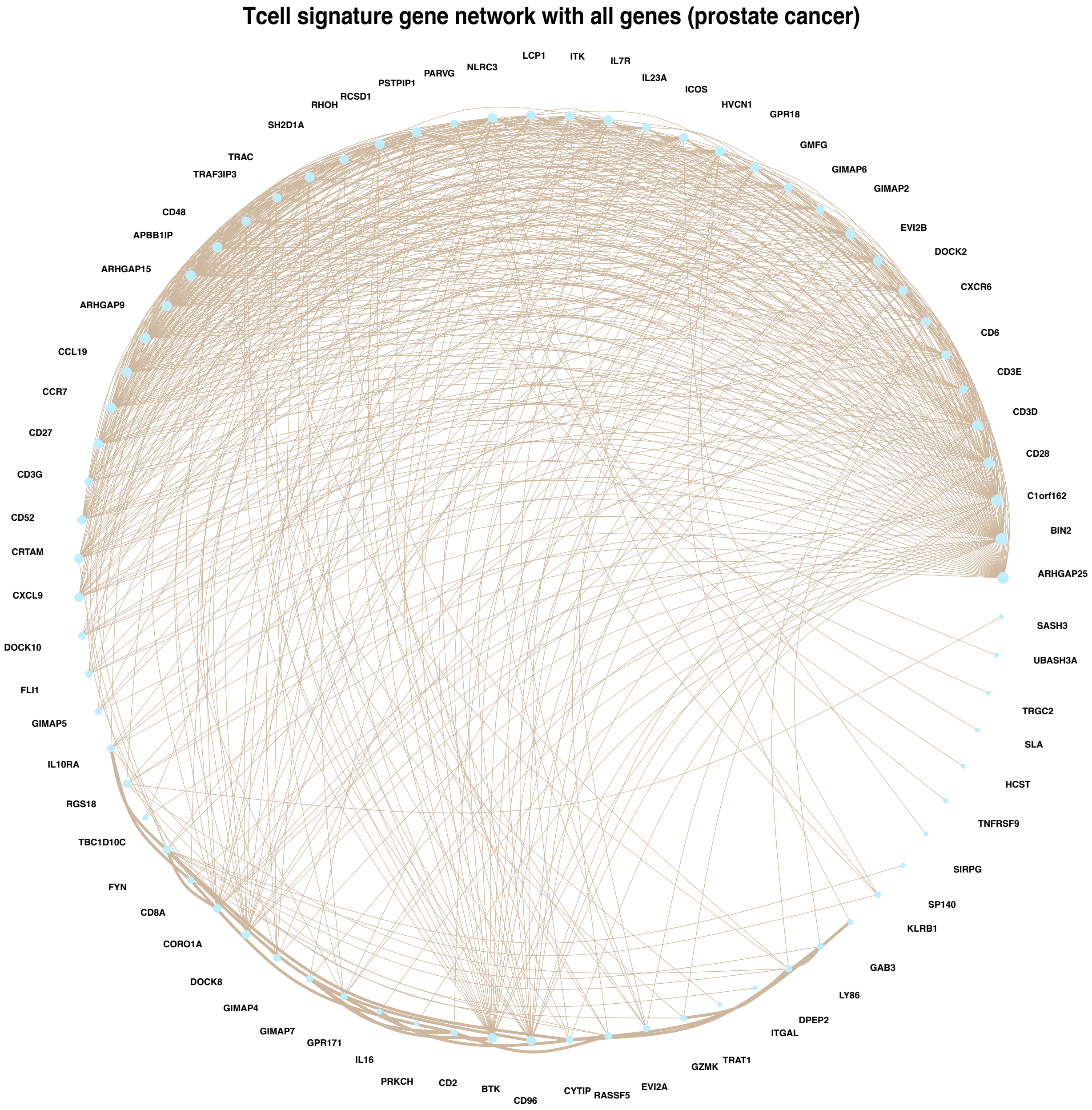


Figure S19: Global spatial connectivity network for T cell immune signature genes where all genes are included for prostate cancer tissue.

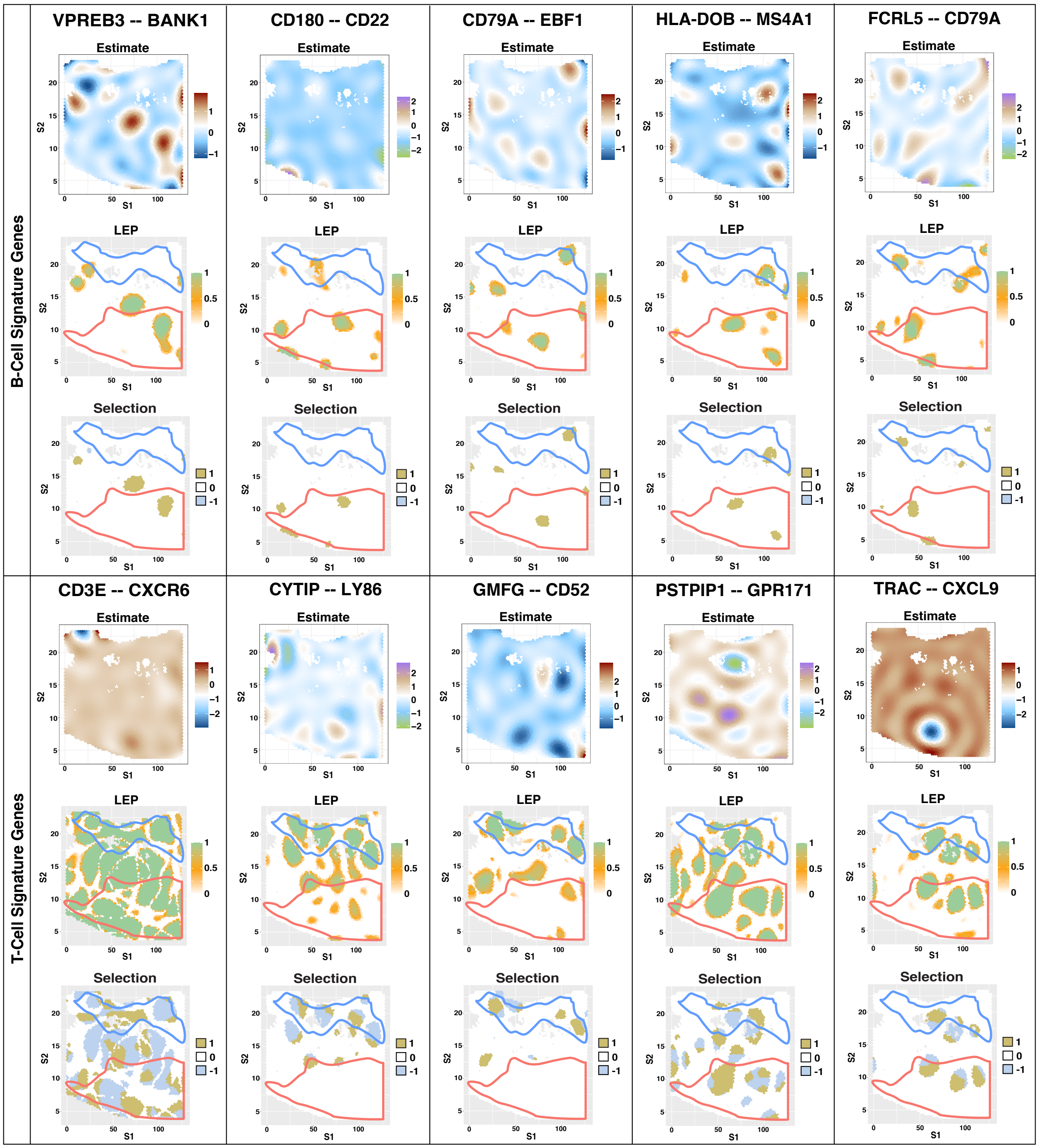


Figure S20: In case of the prostate cancer tissue, the Figure shows spatially varying edges for immune signature genes for B cell (top row: VPREB3 – BANK1, CD180 – CD22, CD79A – EBF1, HLA-DOB – MS4A1, and FCRL5 – CD79A) and T cell (bottom row: CD3E – CXCR6, CYTIP – LY86, GMFG – CD52, PSTPIP1 – GPR171, and TRAC – CXCL9) based on their global posterior inclusion probability. We provide the estimate, spatially varying PIP and selection. The red and blue demarcation on the PIP and selection plot indicates tumor and normal region respectively. Rest of the region is considered as an intermediate region.

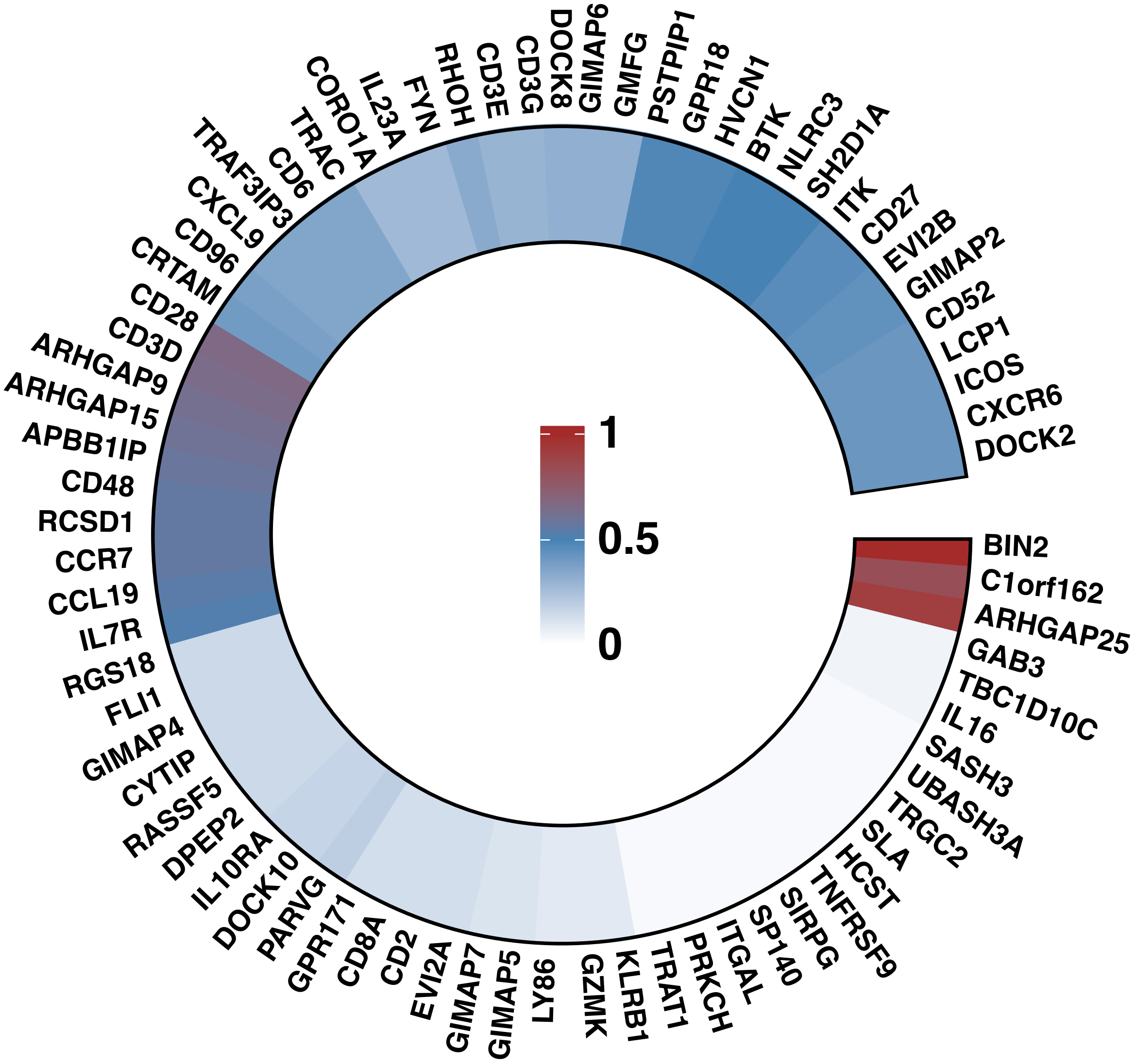


Figure S21: The Figure shows the circular heatmap of a matrix representing the number of connections of a immune signature gene w.r.t. T cell. Here we considered all the Tcell immune signature genes for the prostate cancer tissue.
